# A Mathematical Framework for the Quantitative Analysis of Genetic Buffering

**DOI:** 10.1101/2024.12.09.627635

**Authors:** Jim Karagiannis

## Abstract

Genetic buffering plays a pivotal role in orchestrating the relationship between genotype and phenotype in outbred populations. While high-throughput screens have identified many instances of genetic buffering – through the detection of “synthetic lethality” or “synthetic sickness” – a formal and general method for its quantitative analysis across systems is lacking. In this report, an axiomatic mathematical framework that can be used to classify, quantify, and compare buffering relationships between genes is described. Importantly, this methodology employs a ratio scale as its basis, thereby permitting the definition of a novel neutrality model for gene interaction that is referred to as the “parallel” model. This model does not contradict, and instead complements, the commonly used “product” model (more aptly referred to herein as the “serial” neutrality model). Moreover, simple extensions of this newly developed framework permit the unambiguous definition and classification of gene interactions in a formal, general, and mathematical way. Consequently, the concept of genetic buffering as first conceived by Leland Hartwell becomes a specific case within a comprehensive model of gene interaction.

## Introduction

Predicting the relationship between genotypic variation and phenotypic variation is a fundamental goal of the discipline of genetics (1–6). From the "modern synthesis" that united Darwinian selection and Mendelian genetics in the early part of the 20^th^ century, to the genomic revolution of the early 21^st^ century, the discipline’s inability to reliably delineate the pathway from genotype to phenotype has hampered progress in fields ranging from evolutionary biology to the study of human disease (7–14). Indeed, while major technological leaps forward have been made – primarily through improvements in DNA sequencing together with ever more powerful computational tools (e.g., GWAS, “Omics” technologies, machine learning approaches) – there remains a so-called “genotype-phenotype gap” (4, 15–18). This gap refers to the difficulty encountered in making causal connections between genotypic variation and phenotypic variation and manifests itself most notably in the form of the “missing heritability” problem; a problem borne of the fact that variants identified by GWAS typically explain only a small fraction of a given trait’s heritability (7, 8, 10, 19–21).

One hypothesis put forward to explain this discordant observation posits that the “missing” heritability derives – not from the fact that there are additional variants remaining to be discovered – but instead, from the fact that the heritability estimates themselves fail to incorporate the consequences of gene interaction (22). The discipline’s neglect in appreciating the relevance and importance of gene interaction stems, at least in part, from its historical reliance on inbred model organisms (which in effect limits the genotype-phenotype relationship to the allele of interest only) (23). The observation that alleles often produce distinct phenotypes in different genetic backgrounds highlights the importance of interacting genes (historically referred to as “suppressors”, “enhancers”, or “modifiers”). In yeast, for example, a comparison of Σ1278b and S288C strains (where the genetic divergence is roughly equal to the divergence between two human genomes) reveals that 44 gene deletions are uniquely essential in Σ1278b backgrounds and 13 gene deletions are uniquely essential in S288C backgrounds (24). While the use of inbred organisms improves experimental reproducibility by reducing genetic variation between individuals, this strategy fails to incorporate or consider the genetic diversity that exists in natural, outbred populations. Moreover, it leads to the erroneous impression that the effects of mutation are deterministic in nature.

"Genetic buffering" represents one manner in which gene interactions influence the genotype-phenotype relationship (23, 25–28). The importance of genetic buffering – defined by Hartman as the “compensatory process whereby particular gene activities confer phenotypic stability against genetic or environmental variations” (25) – was first articulated by Nobel laureate Leland Hartwell and colleagues in 2001 (23). At that time Hartwell presciently suggested that buffering relationships between genes might “provide a force to maintain genetic variation in an unexpressed state in some genotypes”. Furthermore, he suggested that such relationships could be revealed by identifying instances of synthetic lethality. The reasoning behind this assertion being that, in these cases, organisms with full function of one of the two genes in question would be able to tolerate variation in the other (i.e., the effects of variation would only be lethal in cases where both gene products had lost activity). In this way, buffering relationships could conceal the phenotypic effects of genetic changes and thereby allow the build-up of “cryptic” variation (28).

In addition to articulating the importance of genetic buffering in influencing the genotype-phenotype relationship, Hartwell also suggested that the comprehensive identification of synthetic lethal interactions in model organisms (via genome-wide screens) might provide a means to study buffering relationships in a systematic and exhaustive manner (23). Since that time, a variety of high-throughput strategies have been developed to identify “synthetic lethal” or “synthetic sick” interactions (synthetic sick interactions being defined as instances where a significant, but non-lethal reduction in fitness is observed) (29–40). In each methodology, the presence and/or strength of an interaction is determined using statistical methods that culminate in the generation of a genetic interaction (GI) score (15, 16, 41–44). Central to this approach is the “neutrality model” chosen to characterize an interaction. Typically, interactions are defined by first assuming that the two mutant alleles (at different loci) act independently. Next, a neutrality model is chosen to predict the combined effect of both mutations under the assumption of independent action. If the double mutant phenotype differs significantly from the expected phenotype under the chosen neutrality model, the two mutant alleles are then said to interact (15, 16, 42).

While the procedure is simple, the choice of neutrality model is equivocal. The multiplicative “product” model (where the expected double mutant phenotype is equal to the product of the fractional effect of each individual mutation) is commonly used. However, “additive”, “min”, and “log” models have also been proposed. Furthermore, the appropriateness of each model as well as the circumstances under which each should be applied remains open to debate (42, 45, 46).

In this report, I propose and describe a previously unrealized methodology for quantifying genetic buffering relationships. This is accomplished through the application of a mathematical framework first put forward by B.M. Schmitt to formally measure buffering action (47, 48). By applying this framework, I develop a novel neutrality model for gene interaction – referred to as the “parallel” neutrality model – that is based on the use of ratio scales, the highest measurement scale possible (49). Importantly, this model does not contradict, and instead complements, the commonly used “product” neutrality model. Moreover, by extending this newly developed framework, I show that it is possible to unambiguously define and compare gene interactions in a formal, general, and mathematical way.

## Results

### Mathematical preliminaries

The first large-scale genetic screen for synthetic sick or synthetic lethal (SSL) interactions was performed in the unicellular model eukaryote, *Saccharomyces cerevisiae* (commonly referred to as “budding” yeast). In this study, colony size – a proxy for fitness – was measured in a set of systematically constructed, haploid, double gene-deletion mutants (39). The approach, referred to as synthetic genetic array analysis (SGA), was later refined and broadened, becoming a fully automated, high-throughput methodology capable of assaying the complete set of both non-essential and essential budding yeast genes (the latter being made possible through the creation of temperature-sensitive alleles). Using the multiplicative “product” neutrality model, the authors identified nearly one million interactions and used this data to create a global genetic interaction map for budding yeast (31).

In these experiments, colony size was used to estimate individual growth rates and hence the fitness of the respective genotypes. Since cellular growth represents a relatively simple phenotype that is easily described mathematically, and furthermore, since it can be used to provide a quantitative determination of fitness, it will be utilized in the paragraphs that follow to illustrate the current methodology. To begin, an exponential growth model will be explored.

Exponential growth can be described by the ordinary differential equation (ODE)

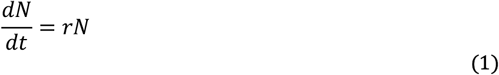

where *N* represents the number of cells in the population and *r* the population growth rate (50). In the context of microbial growth, the parameter, *r*, can be further decomposed and expressed as:

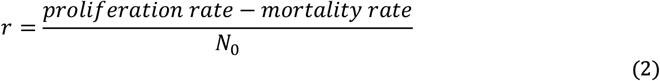

Since fitness in a unicellular organism can be defined by the per capita growth rate, the value of the *r* parameter provides a measure of the fitness of a particular genotype. For example, a genotype with a measured *r* value of 1.4 would be considered to have a 40% fitness advantage over a genotype with a measured *r* value of 1. Similarly, a genotype with an *r* value of 0.5 would be considered to be half as fit as a genotype with an *r* value of 1. Equation (1) can be integrated analytically to give the number of cells as a function of time

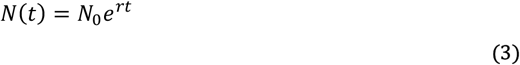

Using the multiplicative “product” neutrality model, it is possible to estimate the expected fitness of a double mutant based on the *r* values of the respective single mutants. To illustrate this, first consider a hypothetical haploid cell type with a measured *r* value of 1/hr (corresponding to a doubling time of 0.693 hours or 41.59 minutes). In addition to this wild-type strain, also consider two other strains: one expressing the mutant ***c***_***1***_ allele, and the other expressing the mutant ***d***_***1***_ allele (where ***c*** and ***d*** define different loci). The growth rate constants (*r*) for the ***c***_***1***_ and ***d***_***1***_ mutants will be assumed to be 0.7/hr and 0.5/hr, respectively. According to the multiplicative “product” neutrality model, the expected fitness of the double mutant, ***c***_***1***_***d***_***1***_, is simply the product of 0.7 and 0.5, or 0.35/hr. Thus, the double mutant would be expected to grow in accordance with the equation, *N(t)=N* _*0*_ *e*^*0*.*35t*^. Thus, if the value of *r* for the double mutant were observed to be significantly less than 0.35/hr, then one would conclude a “negative” interaction exists between the alleles (i.e., the double mutant would be considered to be less fit than expected). In contrast, if the value of *r* for the double mutant were observed to be significantly greater than 0.35/hr, then one would conclude a “positive” interaction exists between the alleles (i.e., the double mutant would be considered to be more fit than expected). Finally, if the value of *r* for the double mutant were not observed to be significantly different from 0.35/hr, then one would conclude that no interaction exists between the alleles (**Figure 1A**).

**Figure 1.**
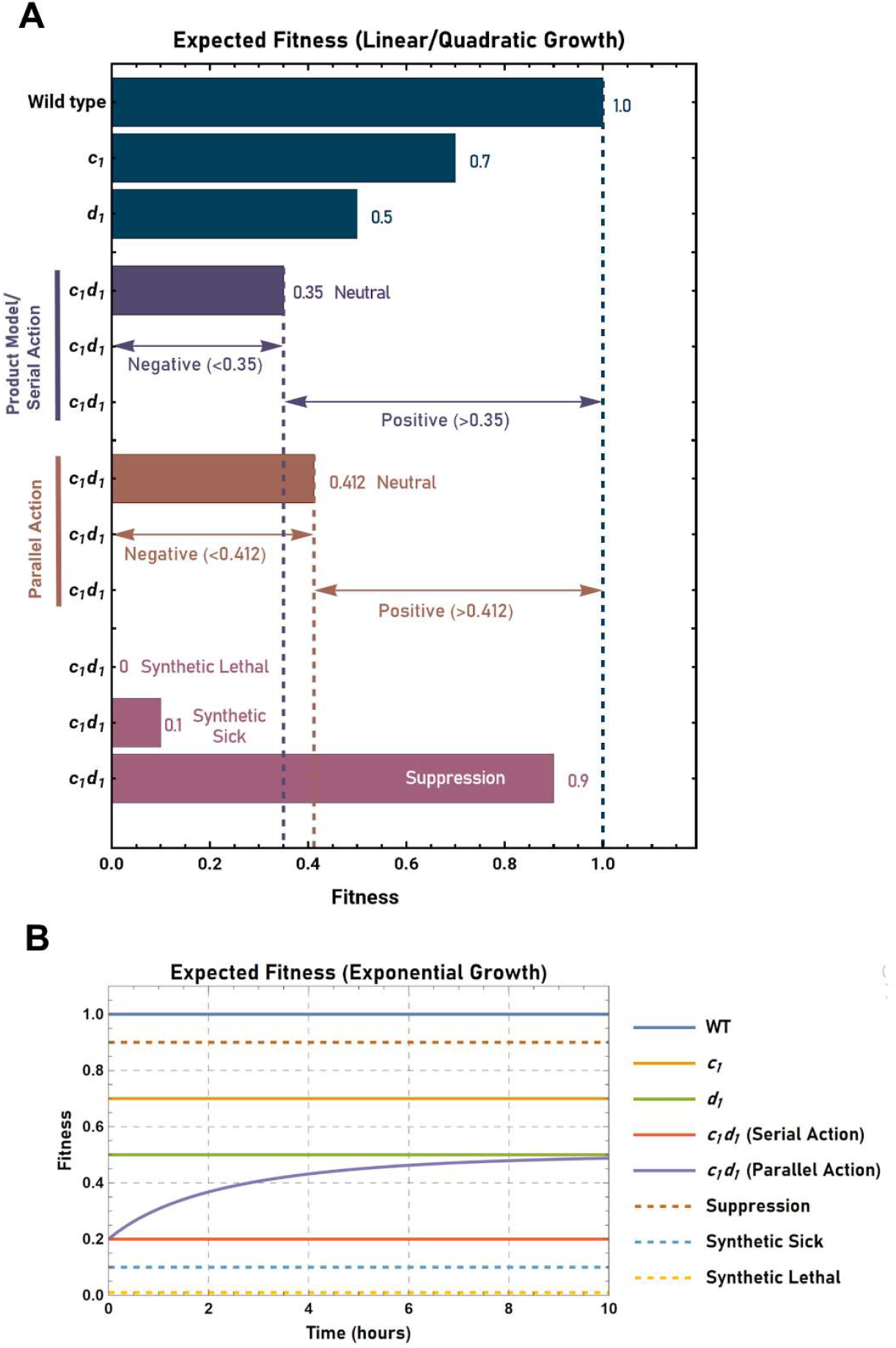
Expected fitness of the *c*_*1*_*d*_*1*_ genotype upon application of either the “serial” or “parallel” neutrality models. **(A)** Expected fitness of the ***c***_***1***_***d***_***1***_ genotype in the context of linear or quadratic growth. In this scenario the “product” neutrality model and the “serial” neutrality model are mathematically equivalent (see text for details). Arrows indicate the range of fitness values corresponding to negative or positive genetic interaction between the ***c***_***1***_ and ***d***_***1***_ alleles for both the product/serial (blue) and “parallel” (brown) neutrality models. Fitness values corresponding to the phenotypes of synthetic lethality, synthetic sickness, or suppression are also shown (magenta). **(B)** Expected fitness of the ***c***_***1***_***d***_***1***_ genotype in the context of exponential growth. Note that in this context the product neutrality model and the serial neutrality model are no longer equivalent. In contrast to serial action, fitness varies as a function of time in scenarios involving parallel acting alleles.

In the sections that follow, the multiplicative “product” neutrality model described above will be contrasted to a novel neutrality model derived from the application of Schmitt’s paradigm for quantifying buffering action (47). Furthermore, the suitability of the product model in the context of exponential growth will be assessed in comparison to its use in the context of linear or quadratic growth. However, prior to comparing these models, a brief explanation of Schmitt’s underlying mathematical framework is necessary.

### Schmitt’s buffering paradigm illustrated using interconnected fluid-filled vessels

Three mathematical functions provide the foundation of Schmitt’s paradigm: the “transfer” function, the “buffer” function, and the “sigma” function. The transfer function describes the observable change in the quantity of interest (i.e., the characteristic of the system being observed and measured) (47). In the case of exponential microbial growth, this quantity is *N*, the number of cells in the population. The buffering function, on the other hand, describes the change in the quantity of interest that <u>would</u> have been observed, if not for the buffering action present in the system. A useful way to visualize such systems (in an abstract and general sense) is to imagine the partitioning of a fluid between two vessels that are connected via a conduit at their base. In this case, a fluid poured into either of the vessels would flow in an unrestricted manner between the two. Thus, the level (i.e., height) of the fluid in each vessel would always be the same (irrespective of the volume of fluid entering). Furthermore, if one poured a fixed volume of fluid into one of the vessels, both the height of the fluid, as well as the volume of fluid in each vessel, would be dependent upon their underlying geometry. One of the two vessels will be referred to as the “transfer” vessel (or “transfer” compartment) and the other as the “buffer” vessel (or “buffer” compartment). Thus, if one unit of fluid is passed into a system consisting of one cylindrical transfer vessel and one cylindrical buffer vessel, and 0.8 units accumulates in the transfer vessel while 0.2 units accumulates in the buffer vessel, then the transfer function would be defined as *τ(x)=0*.*8x* and the buffer function as *β(x)=0*.*2x*. The sigma function describes the quantity of interest that is present in both vessels (in this case the total volume of fluid) and is defined as:

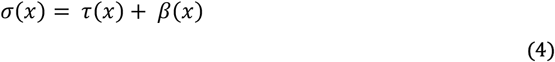

Thus, the sigma function, *σ(x)*, is equal to *x*.

Using this framework, it is now possible to define four buffering parameters: *ŧ, b, T* and *B* (the *ŧ* parameter is indicated with a crossed “*t*” to prevent it being confused with the symbol for time, *t*). The parameter *ŧ* represents the change in the transfer compartment relative to the total change in the system, while the parameter *b* represents the change in the buffer compartment relative to the total change in the system. In contrast, *T* represents the change in the transfer compartment relative to the buffer compartment, and *B* represents the change in the buffer compartment relative to the transfer compartment. Thus, for the system described above, if a further 99 units of fluid were added, a total of 80 units would be present in the transfer compartment and 20 units would be present in the buffer compartment. If the vessels are both assumed to be cylinders (where *Volume=π·radius*^*2*^*·height*) then the radius of the buffer vessel must be one-half that of the transfer vessel. Furthermore, the values of *ŧ, b, T*, and *B* would be 0.8, 0.2, 4, and 0.25, respectively (**Figure 2A**).

**Figure 2.**
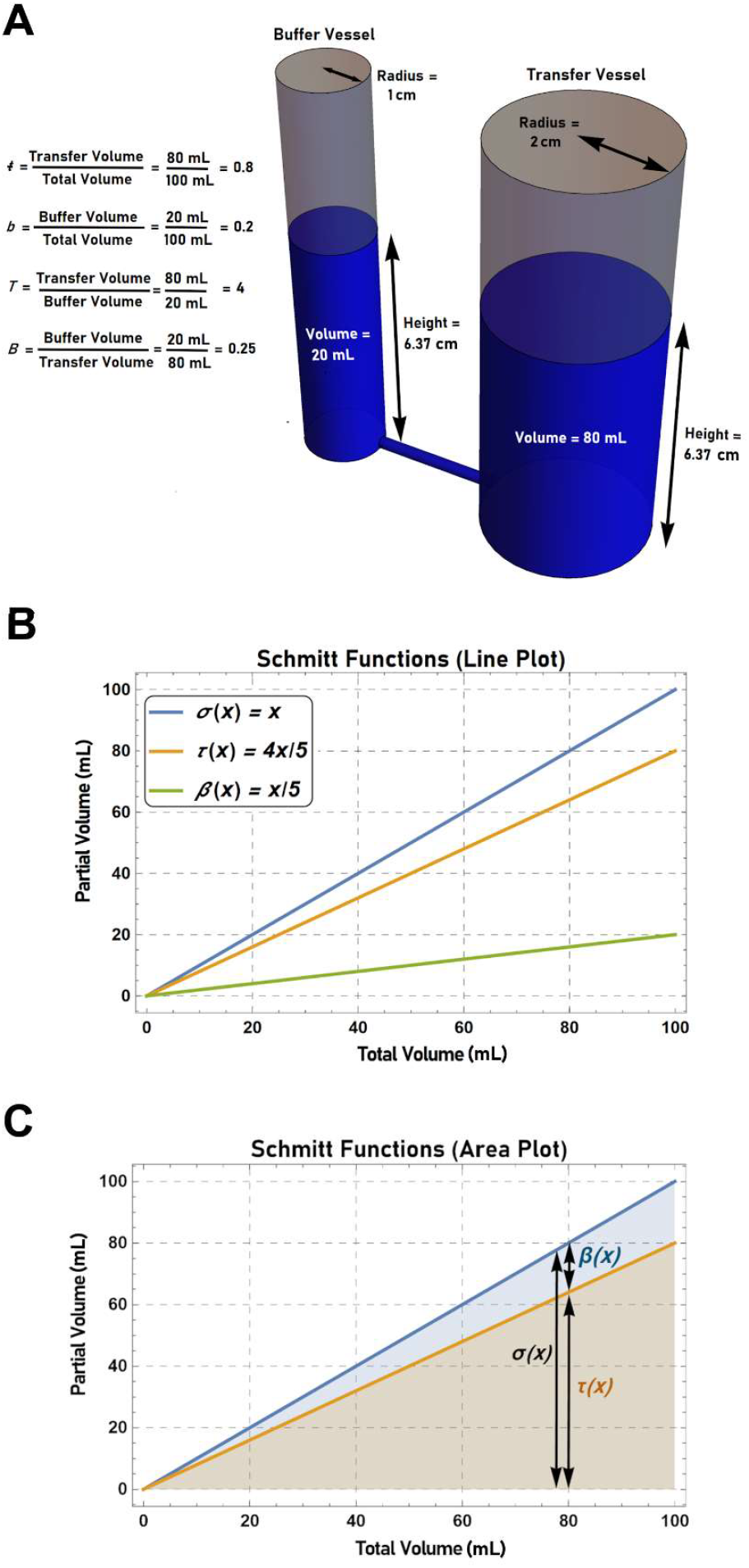
The visualization of buffered systems using interconnected fluid-filled vessels. (**A**) Schmitt’s buffering paradigm (47) can be used to describe any scenario in which there is a partitioning of a measurable quantity between distinct compartments. In this case, 100 mLs of fluid is partitioned between a “transfer” vessel and a “buffer” vessel. Since the two vessels are connected via a conduit at their base, the level (i.e., height) of fluid in each vessel is the same. Furthermore, the amount of fluid that accumulates in each vessel is dependent on its underlying geometry. The effect of the “buffer” vessel on the partitioning of fluid can be described using the *b* parameter (buffer volume/total volume) or the *ŧ* parameter (transfer volume/total volume). Alternatively, the parameters *B* (buffer volume/transfer volume) or *T* (transfer volume/buffer volume) can be used. If a fluid partitions in a ratio of 1:4 between cylindrically shaped transfer and buffer vessels (i.e., *B*=0.25), then the buffer vessel must have a radius one-half that of the transfer vessel. The transfer, buffer, and sigma functions can be plotted as simple line graphs (**B**), or as an area plot (**C**), where the transfer and buffer volumes are indicated by the heights of the buffer area (blue) and transfer area (orange) at any given total volume. The sum of the heights of the buffer and transfer areas at any given value of *x* indicates the total volume described by the sigma function.

While the visualization of fluid filled vessels is useful in developing an intuitive understanding of buffered systems, such scenarios can also be formally described by plotting the transfer, buffer, and sigma functions (**Figure 2B**). One can also use these functions to create area plots where the transfer and buffer volumes are indicated by the heights of the transfer and buffer areas at any given total volume. Similarly, the height of the total area indicates the total volume as described by the sigma function (**Figure 2C**).

Of particular importance to the forthcoming discussion is the *B* parameter (a measure of the amount of fluid that partitions into the buffer vessel relative to the transfer vessel). In a genetical context it will be used to represent the quantity of the measured phenotypic characteristic that remains unexpressed or hidden within an abstract and inferred buffer compartment (as a result of the biological effects of the mutation in question). Similarly, the parameter, *T*, will represent the quantity of the measured phenotypic characteristic that is expressed within a “patent” transfer compartment (i.e., that which is visible to an observer). Importantly, as defined above, the parameters *B* and *T* yield a “ratio” scale (i.e., a scale with a non-arbitrary zero and equal intervals between values) (47). Ratio scales are considered the highest scale possible (49). Lastly, it should be noted that the value of each of the buffering parameters can be calculated if the value of any one of the four parameters is known (**Table 1**).

**Table 1.**
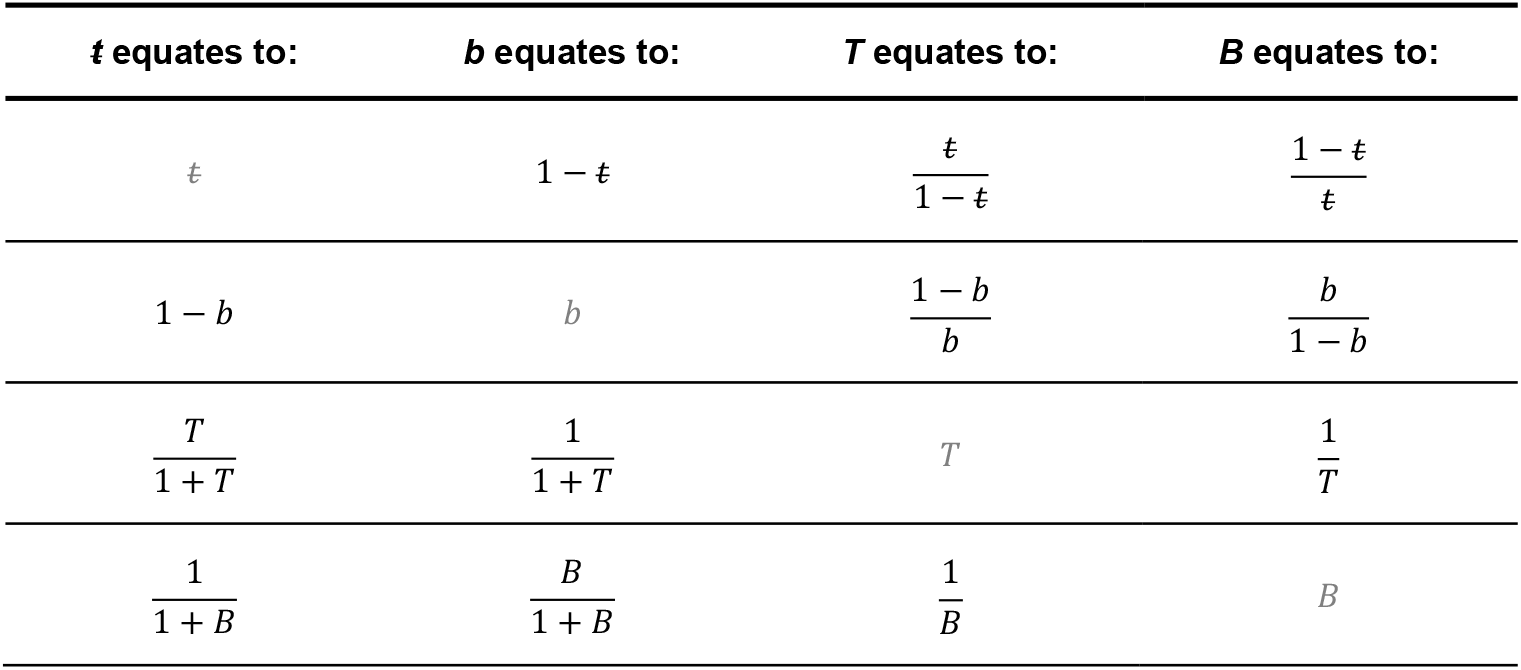
Buffering parameter equivalencies.

### Interactions between vessels

Next, consider the following four systems: **1)** a system consisting of only a cylindrical transfer vessel (i.e., no buffering action is present), **2)** a system consisting of a transfer vessel and a buffer vessel, each identical to the cylindrical transfer vessel in System 1, **3)** a third system identical to the second, and **4)** a system consisting of a transfer vessel identical to that observed in System 1 connected to two buffer vessels that are identical to those observed in Systems 2 and 3 (**Figure 3**).

**Figure 3.**
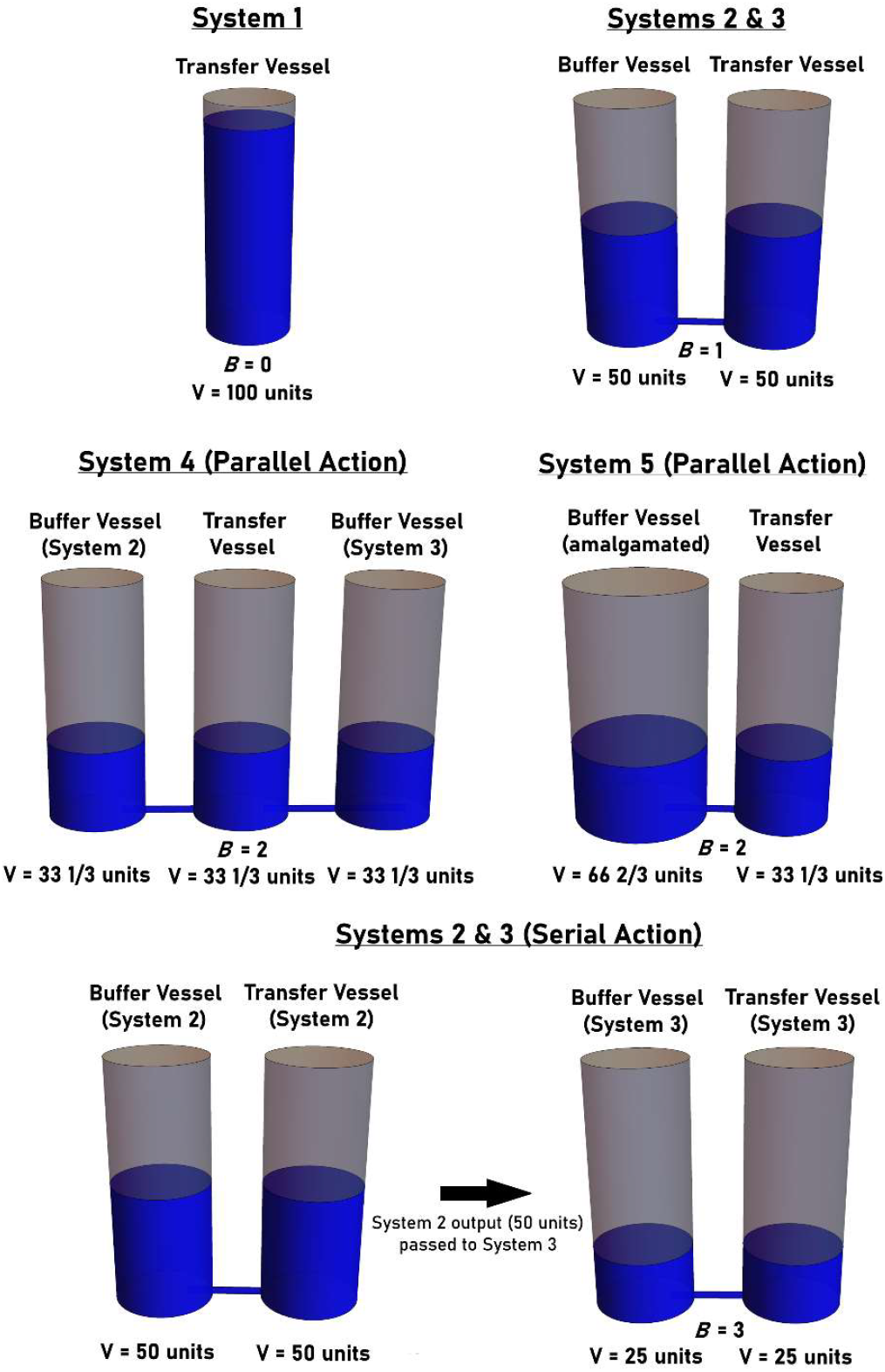
The effects of parallel or serial action on systems of interconnected fluid filled vessels. System 1 (top, left panel) is comprised of only a transfer vessel. Systems 2 and 3 (top, right panel) are comprised of one transfer vessel and one buffer vessel. All buffer and transfer vessels are assumed to be of identical size and geometry. If the buffer vessels act in parallel (i.e., they affect the partitioning of fluid to the transfer vessel simultaneously), their combined effect can be determined by calculating the sum of the *B* values determined for each independently acting vessel (Systems 4 and 5, middle panels). If acting in series (i.e., if the output of System 2 becomes the input to System 3), the combined effect of the buffer vessels can be determined by calculating the product of the *ŧ* values determined for each independently acting vessel (bottom panel).

The buffering parameters for System 1 can be calculated to be: *ŧ* = 1, *b* =0, *B*=0, *T* is undefined (due to division by zero). The buffering parameters for Systems 2 and 3, on the other hand, can be calculated to be: *ŧ* = 0.5, *b* =0. 5, *T*=1, *B*=1. The critical question then becomes: What are the values of the buffering parameters of System 4? Since *B* yields a ratio scale, the value of *B* in System 4 can be calculated by simply determining the sum of the individual *B* values calculated for Systems 2 and 3 (assuming of course that the shape and volume of the transfer vessel remains constant for all four systems). Thus, *B=2* in System 4 (i.e., the fluid partitions in a ratio of 2:1 in the buffer compartments relative to the transfer compartment). This is intuitive since the combined effect of the two individual buffer vessels would be the same as that observed in an alternate system (System 5) where the two buffer vessels were replaced with a single “merged” buffer vessel (i.e., a vessel that was made by the amalgamation of the individual buffer vessels from Systems 2 and 3) (**Figure 3, middle panels**). Next, since the value of any of the buffering parameters can be calculated provided that the value of one of the other parameters is known, it is possible to determine that for System 4: *ŧ* = 1/3, *b*=2/3, and *T*=0. 5 (i.e., the fluid partitions in a ratio of 1:2 in the transfer compartment relative to the buffer compartments). This result is also intuitive, since, if one observed 100 units of a fluid partitioned into three identical, interconnected vessels, then the volume of the fluid in each vessel would be 33 1/3 units.

It is critical to note that in this scenario, the volume in the transfer vessel of System 4 is not calculated by simply multiplying the total volume of fluid by the individual fractional effects of the buffering compartments (i.e., by multiplying 0.5 by 0.5 by 100 units). In fact, a final volume of 25 units could only be achieved by first pouring 100 units of fluid through System 2, taking the fluid that accumulated in the transfer compartment (50 units), and then pouring this volume into System 3 (leaving 25 units of fluid in both the buffer and transfer compartments). In other words, a final transfer volume of 25 units could only be achieved if the output of System 2 became the input to System 3 (i.e., if the fluid was passed <u>sequentially</u> through System 2 and then through System 3) (**Figure 3, bottom panels**). Thus, when determining the combined effect of two independently acting buffering vessels, one must first consider whether the vessels act in parallel or in series. When acting in parallel the final state of the system can be determined by calculating the sum of the *B* values of the individual buffering vessels. In contrast, when acting in series the final state of the system can be determined by calculating the product of the *t* values of the individual buffering vessels.

### Applying Schmitt’s paradigm to linear, quadratic, and exponential growth

To further illustrate the utility of the current methodology, and to demonstrate why the use of the product model in the context of exponential growth is logically flawed, it is necessary to first consider both linear and quadratic growth models (where the product model is indeed suitable) and contrast them to exponential growth. For the sake of simplicity, the increase in volume (*V*) of fluid in a transfer vessel over time (*t*) will be considered using three distinct growth models: **1)** a linear model described by the function *V*(*t*) = *V*_O_*rt*, **2)** a quadratic model described by the function *V*(*t*) = *V*_O_*rt*^2^, and **3)** an exponential growth model described by the function *V*(*t*) = *V*_O_*e*^rt^. The value of *V*_*0*_ will be assumed to be 1 (again for the sake of simplicity).

For each growth model, four systems will be considered: **(i)** a system consisting of only a transfer vessel, where *r* =1, **(ii)** a system consisting of an identical transfer vessel linked to a buffer vessel, where *r* =0. 7, **(iii)** a system consisting of an identical transfer vessel linked to a buffer vessel, where *r* =0 .5, and **(iv)** a system consisting of an identical transfer vessel linked to buffer vessels identical to those in systems (ii) and (iii), and for which the value of the *r* parameter is to be determined. Lastly, each set of systems will be considered in the case of either serial or parallel action.

### Linear growth in the context of serial action

In the case of linear growth, *V(t)* for each system can be described by the functions:

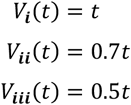

Thus, the Schmitt functions for system (ii) are:

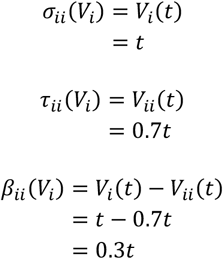

and the Schmitt functions for system (iii) are:

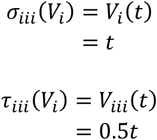

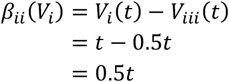

Assuming serial action, the value of the *t* parameter for system (iv) is:

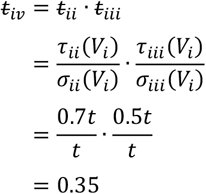

Finally, it is possible to solve for *r*_*iv*_ as follows:

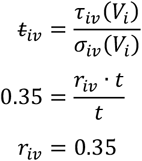

Thus, as expected, *r*_*iv*_ is a constant that is equal to the product of the respective *r* values of systems (ii) and (iii).

### Quadratic growth in the context of serial action

In the case of quadratic growth, *V(t)* for each system can be described by the functions:

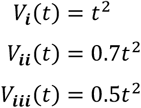

Thus, the Schmitt functions for system (ii) are:

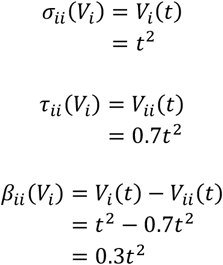

and the Schmitt functions for system (iii) are:

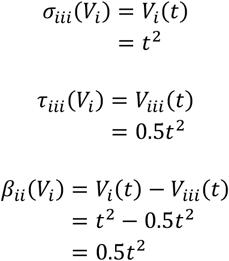

Assuming serial action, the value of the *ŧ* parameter for system (iv) is:

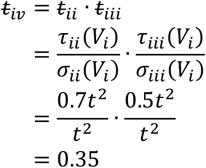

Finally, it is possible to solve for *r*_*iv*_ as follows:

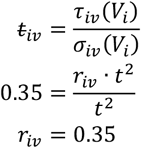

Thus, again as expected, *r*_*iv*_ is a constant that is equal to the product of the respective *r* values of systems (ii) and (iii). Next, the question of whether the same is true in the case of exponential growth will be explored.

### Exponential growth in the context of serial action

In the case of exponential growth, *V(t)* for each system can be described by the functions:

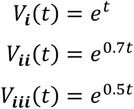

Thus, the Schmitt functions for system (ii) are:

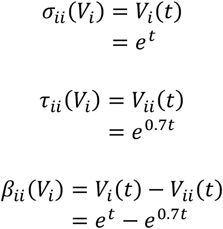

and the Schmitt functions for system (iii) are:

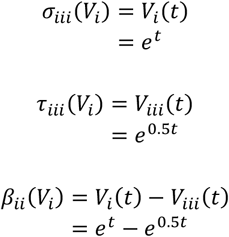

Assuming serial action, the value of the *ŧ* parameter for system (iv) is:

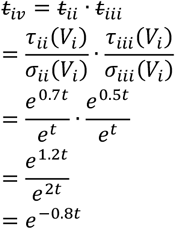

Finally, it is possible to solve for *r*_*iv*_ as follows:

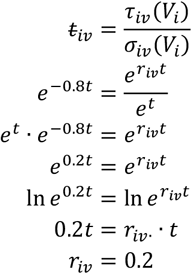

Therefore, in this scenario *r*_*iv*_ is indeed a constant, but is not calculated by determining the product of the respective *r* values of systems (ii) and (iii). Instead, *r*_*iv*_ is calculated by first determining the product of the respective *t* values of systems (ii) and (iii) and then solving for *r*. Thus, in the scenario described above, *r*_*iv*_ can be calculated according to the formula:

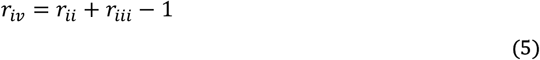

### Linear growth in the context of parallel action

When considering linear growth in the context of parallel action, the equations describing *V(t)* and the Schmitt functions are identical to those described for linear growth in the context of serial action. However, under the assumption of parallel action, the combined effect of both buffer vessels is calculated by determining the sum of the respective *B* parameters. The value of the *B* parameter for system (iv) is:

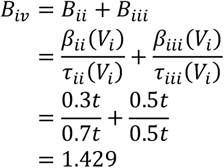

Thus, *ŧ*_*iv*_ can be calculated as follows:

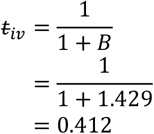

Finally, it is possible to solve for *r*_*iv*_ as follows:

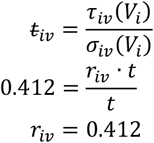

Thus, as expected, *r*_*iv*_ is a constant that can be derived through first determining the sum of the respective *B* values of systems (ii) and (iii), then determining *t*_*iv*_, and then finally solving for *r*_*iv*_.

### Quadratic growth in the context of parallel action

In this scenario the equations describing *V(t)* and the Schmitt functions are identical to those described for quadratic growth in the context of serial action. However, under the assumption of parallel action, the combined independent effect of both buffer vessels is calculated by determining the sum of the respective *B* parameters. The value of the *B* parameter for system (iv) is:

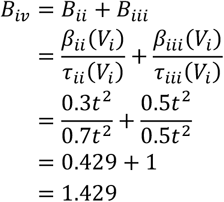

Thus, *ŧ*_*iv*_ can be calculated as follows:

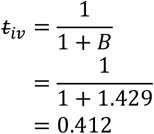

Finally, it is possible to solve for *r*_*iv*_ as follows:

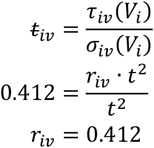

Thus, as expected, *r*_*iv*_ is a constant that can be derived through first determining the sum of the respective *B* values of systems (ii) and (iii), then determining *t*_*iv*_, and then finally solving for *r*_*iv*_.

### Exponential growth in the context of parallel action

In this scenario the equations describing *V(t)* and the Schmitt functions are identical to those described for exponential growth in the context of serial action. However, under the assumption of parallel action, the combined effect of both buffer vessels is calculated by determining the sum of the respective *B* parameters. The value of the *B* parameter for system (iv) is:

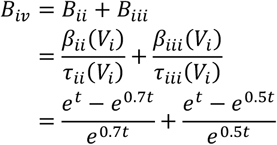

Thus, *ŧ*_*iv*_ can be calculated as follows:

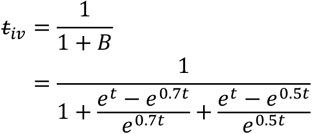

Finally, it is possible to solve for *r*_*iv*_:

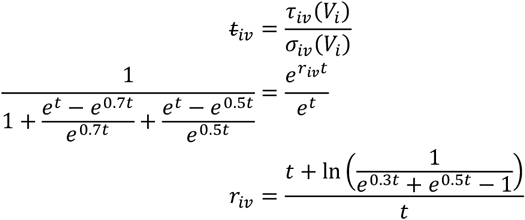

Thus, unlike the previous two scenarios, *r*_*iv*_ is not a constant. However, similar to the previous two scenarios, *r*_*iv*_ is derived through first determining the sum of the respective *B* values of systems (ii) and (iii), then determining *t*_*iv*_, and then finally solving for *r*_*iv*_.

The implications of these results with respect to the phenotype of cellular growth are explored in the section below.

### Applying Schmitt’s paradigm to cellular growth

In the same way it was used to describe systems of fluid-filled vessels, Schmitt’s paradigm can also be used to describe other scenarios in which there is a partitioning of a measurable quantity into distinct compartments. Indeed, the mathematical description of exponential microbial growth, *N(t)=N*_*0*_ *e*^*rt*^, is identical to that used to describe the exponential increase in the volume of fluid accumulating within a transfer vessel, *V(t)=V*_*0*_ *e*^*rt*^. Thus, in applying Schmitt’s paradigm to examples of linear, quadratic, and exponential cellular proliferation, it becomes clear that the product model is inadequate in terms of assessing fitness across distinct modes of growth.

To expound upon these conclusions, consider again the scenario described earlier involving a hypothetical haploid cell-type (where *r*=1/hr) as well as two derived mutant strains expressing either the ***c***_***1***_ or ***d***_***1***_ alleles (where *r* =0.7/hr and *r* =0.5/hr, respectively). Using the generality of Schmit’s paradigm it is now clear that two alternative scenarios must be considered when dealing with two independently acting alleles affecting growth: one in which the two alleles act in <u>series</u> (i.e., the biological effect of the ***c***_***1***_ allele is required for the biological effect of the ***d***_***1***_ allele, or vice versa) and a second in which the two alleles act in <u>parallel</u> (i.e., the biological effect of the ***c***_***1***_ allele <u>is not</u> required for biological effect of the ***d***_***1***_ allele and vice versa). Furthermore, it is important to note that it is impossible to know *a priori* which model to apply in order to assess whether an interaction between two alleles exists.

Nevertheless, in either scenario, the buffering effect of the ***c***_***1***_ allele on exponential growth can be described by the buffering parameters as follows: *t=e*^*-0*.*3t*^, *b*=*1 - e*^*-0*.*3t*^, *T* = *e*^*0*.*7t*^*/(e*^*t*^ *- e*^*0*.*7t*^*), B*= *(e*^*t*^*- e*^*0*.*7t*^*)/e*^*0*.*7t*^. In other words, since the environment can support a growth rate of 1/hr in a wild-type strain, the expression of the ***c***_***1***_ allele results in 1 -*e*^*-0*.*3t*^ percent of this growth potential being “lost” or “diverted” into an abstract buffer compartment. Similarly, the buffering effect of the ***d***_***1***_ allele can be described by the buffering parameters as follows: *t=e*^*-0*.*5t*^, *b*=*1 -e*^*-0*.*5t*^, *T*= *e*^*0*.*5t*^*/(e*^*t*^ *- e*^*0*.*7t*^*), B* =*(e*^*t*^ *- e*^*0*.*5t*^*)/e*^*0*.*5t*^. Using similar logic, the expression of the ***d***_***1***_ allele results in 1 -*e*^*-0*.*5t*^ percent of the wild-type growth potential being lost to an abstract buffer compartment.

With this information, it is now possible to directly and unambiguously calculate the predicted effect of both mutations on exponential growth under the assumption the alleles act independently. Assuming serial action, the expected value of the *ŧ* buffering parameter is *ŧ =e*^*-0*.*8t*^ and the double mutant is expected to grow in accordance with the equation:

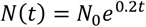

Assuming parallel action, the expected value of the *B* buffering parameter is:

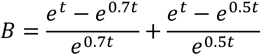

and the double mutant is expected to grow in accordance with the equation:

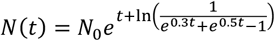

In applying Schmitt’s paradigm, it thus becomes clear that two potential sources of error exist upon application of the product model. First, the product model neglects to consider whether the alleles in question act sequentially or in parallel. Second, the application of the product model in the context of exponential growth (with serially acting alleles) results in an erroneous calculation of the expected double mutant fitness. The difference in fitness calculated by applying the additive “parallel” model versus the multiplicative “product” model in the context of linear or quadratic growth is shown using a heatmap in **Figure 4A**. A second heatmap describing the difference in fitness calculated by applying the “serial” model versus the product model in the context of exponential growth is shown in **Figure 4B**.

**Figure 4.**
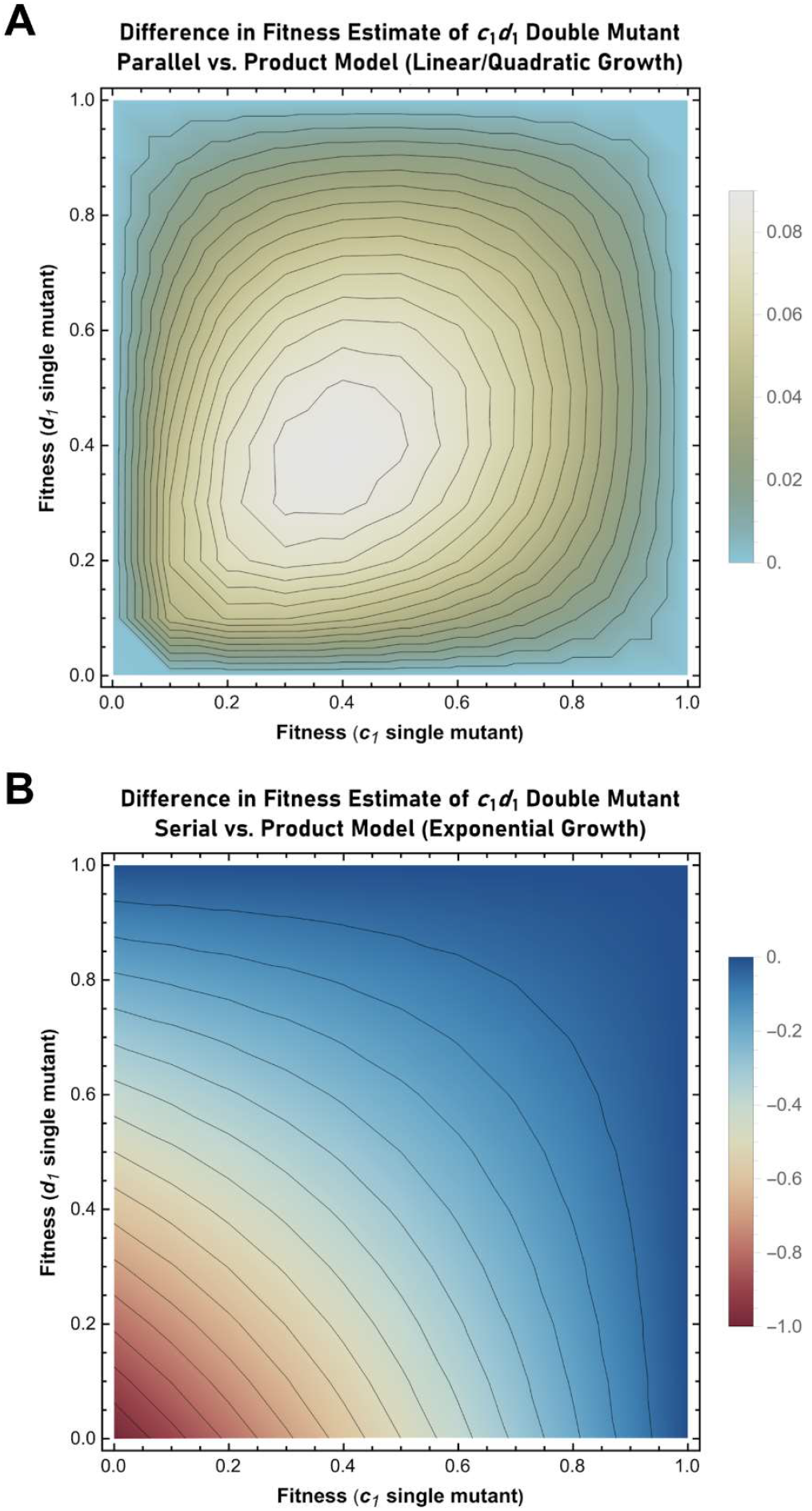
Heat maps describing the difference in fitness (*f*) estimated for the *c*_*1*_*d*_*1*_ genotype upon application of either the parallel or serial neutrality models in place of the product model. **(A)** Difference in the estimated fitness of the ***c***_***1***_***d***_***1***_ genotype when the “parallel” model is applied in place of the “product” model (in the context of linear/quadratic growth). *f*_*parallel*_ is determined by calculating the sum of the individual *B* parameters that describe the respective ***c***_***1***_ and ***d***_***1***_ single mutants and then solving for *r* (see text for details). *f*_*product*_ is determined by calculating the product of the individual *r* values that describe the respective ***c***_***1***_ and ***d***_***1***_ single mutants. The heat map displays the calculated value of *f*_*parallel*_ – *f*_*product*_ for mutant alleles ranging in fitness from 0.01 to 1. **(B)** Difference in the estimated fitness of the ***c***_***1***_***d***_***1***_ genotype when the “serial” model is applied in place of the “product” model (in the context of exponential growth). *f*_*serial*_ is determined by calculating the product of the individual *ŧ* parameters that describe the respective ***c***_***1***_ and ***d***_***1***_ single mutants and then solving for *r* (see text for details). *f*_*product*_ is determined by calculating the product of the individual *r* values that describe the respective ***c***_***1***_ and ***d***_***1***_ single mutants. The heat map displays the calculated value of *f*_*serial*_ – *f*_*product*_ for mutant alleles ranging in fitness from 0 to 1.

To avoid confusion moving forward, the application of Schmitt’s paradigm to determine the expected fitness of double mutants in the context of either serial or parallel action will hereafter be referred to as:

1) Theequations/627635v1 “serial” neutrality model. This model is identical to the multiplicative “product” model in the context of linear or quadratic growth, but not in the context of exponential growth (**Figure 1A, B**). Using this model, the expected phenotype (visualized as the volume partitioned to the final transfer compartment) can be calculated by determining the product of the *t* buffering parameters associated with the independent action of two (or more) alleles functioning in a sequential manner.
2) The “parallel” neutrality model. This model is distinct from the product model in the contexts of linear, quadratic, or exponential growth (**Figure 1A, B**). In this case, the additivity of the *B* parameter can be used to determine the total growth that is buffered due to the independent parallel action of the alleles (visualized as the volume partitioned to the buffering compartments of the system in question). The expected growth (visualized as the volume partitioned to the transfer compartment relative to the total volume) can then be determined using the formula, *t* = 1/(1 +*B*). Lastly, it should be noted that the expected combined effect of any combination of two (or more) independently acting alleles functioning in parallel can be calculated by first evaluating the sum of their *B* values and then determining the corresponding *t* value.

### The buffering effect of individual alleles versus genetic buffering

Using the methodology presented above, it is possible to calculate the “buffering” effect of any allele, as well as the expected combined effect of two or more alleles using either the “parallel” or “serial” neutrality models. At this juncture it is necessary to make clear that the use of the term “buffering” in this context is distinct from the term “genetic buffering” as proposed by Hartwell. The former pertains to the determination of the phenotypic effect of individual alleles via Schmitt’s paradigm, while the latter pertains to an interaction between alleles leading to synthetic lethality. This conflation of terms is unfortunate, but nevertheless, any confusion can be easily resolved by precisely defining the individual concepts mathematically using the current framework.

The buffering effect of any individual allele on growth is defined by the parameter, *B*, on a scale that ranges from zero, indicating no buffering/perfect transfer (i.e., the allele has no effect on growth), to infinity. In scenarios where there is perfect buffering/no transfer (i.e., the allele prevents any growth from occurring) the value of *B* becomes undefined due to division by zero. In contrast, genetic buffering as conceived by Hartwell pertains to a relationship between two loci, where:

1) in backgrounds where the allele at the first locus is fully functional, expression of a non-functional allele at the second locus results in an insignificant deviation of *B* away from zero; and
2) in backgrounds where the allele at the second locus is fully functional, expression of a non-functional allele at the first locus results in an insignificant deviation of *B* away from zero; and
3) expression of non-functional alleles at both loci results in *B* approaching infinity relative to a reference strain expressing fully functional alleles at both loci.

In other words, Hartwell’s concept of genetic buffering can be defined using Schmitt’s paradigm as a specific case in which two individual alleles at two distinct loci – each exhibiting *B* values near zero relative to a wild-type reference strain – unexpectedly combine to result in a double mutant strain that exhibits a *B* value approaching infinity (i.e., when *B*_*observed*_ >> *B*_*expected*_). Using this framework, synthetic sickness is simply the sub-set of such negative interactions where the growth rate of the double mutant is observably greater than zero. Synthetic lethality, on the other hand, is the sub-set of such negative interactions where the growth rate of the double mutant is not observably greater than zero.

It is important to note that in such cases, cells expressing a fully functional allele at the first locus are free to accumulate genotypic variation at the second locus without that variation producing selectable phenotypic variation (at least with respect to growth). It should also be noted that such “cryptic” genotypic variation might be revealed as selectable phenotypic variation by further mutations that result in the expression of loss of function alleles at the first locus of the interacting pair (and vice-versa).

In addition, as should now be intuitively apparent, it is possible to quantify the potential for genetic buffering using the aid of the *B* parameter. For example, the potential for genetic buffering is maximized in the case of a synthetically lethal allele pair where each individual allele exhibits a *B* value near zero (i.e., when *B*_expected_ is also near zero and *B*_observed_ approaches infinity). In such cases, allelic variation at either locus would accumulate in the complete absence of any associated phenotypic variation provided the allele at the other locus was fully functional (thereby maximizing the potential for cryptic genotypic variation). In contrast, the potential for genetic buffering approaches a minimum in the case of a synthetically lethal allele pair where each individual allele exhibits a *B* value (relative to a wild-type reference) that is itself approaching infinity. In this scenario, genotypic variation becomes completely unconstrained with respect to creating phenotypic variation (i.e., cryptic genotypic variation is not possible). In intermediate cases, genetic buffering is possible, but genotypic variation is quantifiably constrained with respect to creating phenotypic variation (i.e., the potential for cryptic genotypic variation is present, but not maximized). Thus, using the current framework, Hartwell’s original qualitative description of genetic buffering (23) becomes fully quantitative.

To fully explore the utility of the current paradigm in unambiguously quantifying genetic buffering, and to do so in both an intuitive and easily visualizable manner, it is necessary to introduce a fifth buffering parameter referred to as the buffering angle, *α*. The buffering angle permits the representation of any form of buffering without the introduction of discontinuities. Such discontinuities appear in the cases of perfect buffering or perfect transfer (where the parameters *B* and *T* are respectively undefined due to division by zero). The advantages of using the buffering angle to quantify genetic buffering is examined in detail below.

### Quantifying the buffering effects of individual alleles using the buffering angle

As described by Schmitt (47), the utility of the buffering angle, *α*, is derived from its capacity to express the buffering properties of a system using a single-value measure – an angle ranging from -45° to 135° – and to do so without the introduction of discontinuities. These discontinuities arise due to division by zero in systems exhibiting perfect transfer (where *T* is undefined since the quantity partitioned to the buffering compartment is zero) and in systems exhibiting perfect buffering (where *B* is undefined since the quantity partitioned to the transfer compartment is zero).

The buffering angle is defined by plotting *ŧ* on the horizontal X-axis (abscissa) and *b* on the vertical Y-axis (ordinate) and determining the angle created between the X-axis and the line drawn from the origin (0,0) to the point defined by (*ŧ*,*b*). In the case of perfect buffering (*ŧ* = 0, *b* =1), the angle between this line – the line joining (0,0) and (0,1) – corresponds to 90°. In the case of perfect transfer (*ŧ* = 1, *b*=0), the angle between the X-axis and the line joining (0,0) to (1,0) corresponds to 0°. In the case of equal transfer (*ŧ* = 0.5, *b*=0. 5) the angle between the X-axis and the line joining (0,0) to (0.5,0.5) corresponds to 45° (**Figure 5**). Thus, the buffering properties of a system in which the values of *ŧ* and *b* fall between 0 and 1 can be described with an angle ranging from 0° to 90° (corresponding to values of the *B* parameter ranging from zero to +∞ and becoming undefined at an angle of exactly 90°).

**Figure 5.**
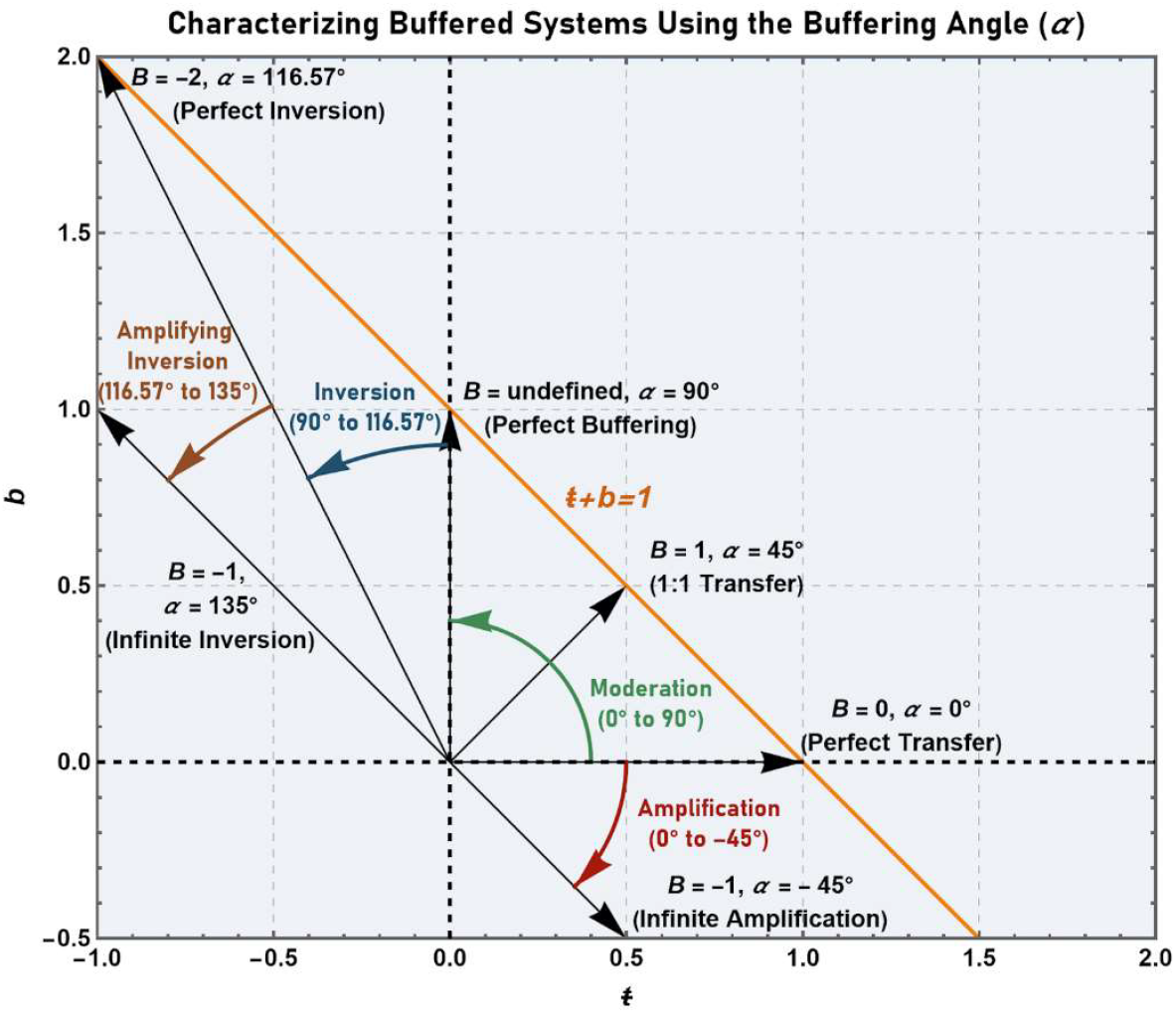
Using the buffering angle to classify and characterize buffered systems. The buffering angle is defined by plotting the line, *b*=1-*ŧ*, and determining the angle between the *ŧ* -axis and the line joining the origin (0,0) to the point (*ŧ, b*). Perfect buffering is thus represented by an angle of 90° and perfect transfer by an angle of 0°. Intermediate angles (i.e., those between 0° and 90°) indicate “moderating” buffered systems. Angles ranging from 0° to -45° (corresponding to *ŧ* values greater than 1 and *b* values less than 0) indicate “amplifying” buffered systems. Angles ranging from 90° to 116.57° (corresponding to *ŧ* values less than zero and *b* values greater than 1) indicate “inverting” buffered systems. Finally, angles ranging from 116.57° to 135° indicate buffered systems that are both “inverting” and “amplifying".

As defined above, this scale (where the sum of *ŧ* and *b* must be equal to 1) is also able to accommodate negative *b* or *ŧ* values. In the context of growth, *ŧ* values greater than one (meaning *b* values are negative) represent cases where the value of the growth constant *r* is greater than that observed in the wild-type reference strain. Thus, angles ranging from 0° to - 45° represent an “amplification” of the quantity (phenotype) of interest and correspond to *B* values ranging from 0 to -1. Values of the *ŧ* parameter that are less than zero (corresponding to *b* values that are greater than one) represent cases where the quantity (phenotype) of interest becomes inverted. To conceptualize such a scenario, it is only necessary to consider the situation where the mortality rate exceeds the proliferation rate, thereby resulting in negative growth (and therefore a negative *ŧ* value). Thus, angles ranging from 90° to 116.57° (corresponding to *ŧ* values ranging from 0 to -1) represent an “inversion” of the phenotype. Such scenarios are described by *B* values ranging from undefined to -2.

Values of the *ŧ* parameter less than -1 (corresponding to *b* values that are greater than two) represent cases where the quantity of interest in the mutant is not only inverted, but inverted to such a degree that the magnitude of negative transfer (observed in the mutant) exceeds the magnitude of transfer in the positive direction (observed in the wild-type reference strain). This corresponds to the expression of alleles where the absolute value of the mortality rate exceeds the absolute value of the proliferation rate. Such scenarios could be described with angles ranging from 116.57° to 135° and with *B* values ranging from -2 to -1. In this way the buffering effect of any allele – even abstract scenarios (that may or may not be physically possible) – can be described by a single finite angle between -45° and 135°. A schematic summarizing the interpretation of the buffering angle, *α*, is shown in **Figure 5**.

It is also of interest to note that the buffering angle can be defined by plotting a three-dimensional space curve of the system, where the total volume is represented on the X-axis (abscissa), the transfer volume on the Y-axis (ordinate), and the buffer volume on the Z-axis (applicate) (**Figure 6A, B**). Interestingly, a projection of this curve onto the X-Y plane produces the transfer function (i.e., *τ(x)* vs. *σ(x)*), whereas a projection of the same curve onto the X-Z plane produces the buffer function (i.e., *β(x)* vs. *σ(x)*) (**Figure 6C, D**). Lastly, the projection of the curve onto the Y-Z plane produces a function that describes the partitioning of the quantity of interest in the buffer vessel versus the transfer vessel (i.e., *β(x)* vs. *τ(x)*). In this projection, the angle between the Y-axis and a line defined by joining the origin to the coordinates (*τ(x), β(x)*) defines the buffering angle. Similarly, the angle between the Y-plane and a line defined by joining the origin to the coordinates (*σ(x), τ(x), β(x)*) defines the buffering angle in the 3D space curve (**Figure 6E**).

**Figure 6.**
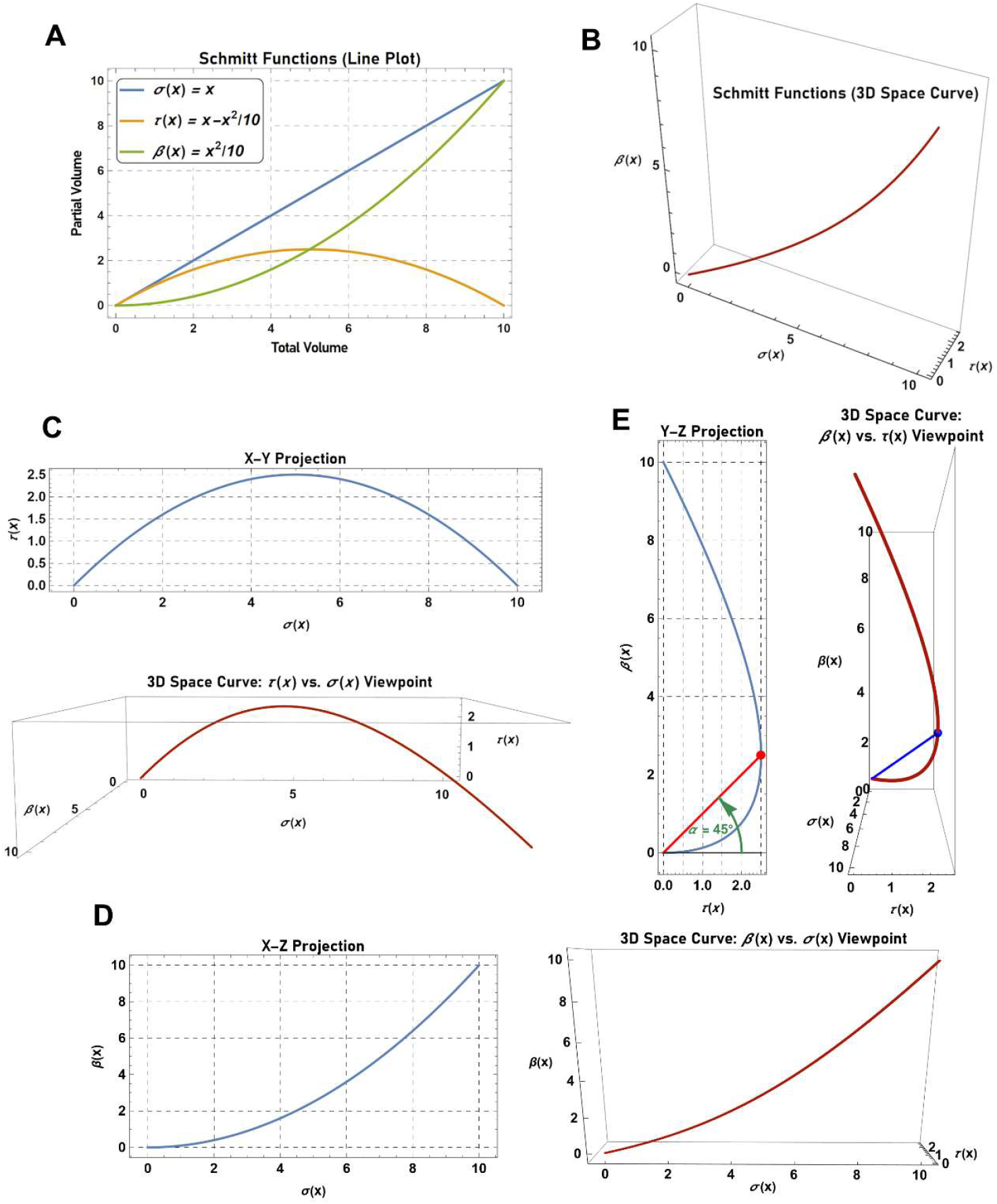
Using three-dimensional (3D) space curves to graphically describe buffered systems. (A) Line plot describing a buffered system defined by the functions, *σ*(*x*) = *x, τ*(*x*) = *x*-*x*^2^-10, and *β* (*x*) =*x*^2^/ 10. (B) 3D space curve of the system defined in (A). The curve is created by plotting *σ*(*x*) on the X-axis (abcsissa), *τ* (*x*) on the Y-axis (ordinate), and *β*(*x*) on the Z-axis (applicate). (C) The X-Y projection of the 3D space curve produces a plot of *τ*(*x*) vs. *σ*(*x*). (D) The X-Z projection of the 3D space curve produces a plot of *β*(*x*) vs. *σ*(*x*). (E) The Y-Z projection of the 3D space curve produces a plot of *β*(*x*) vs. *τ*(*x*). In the Y-Z projection, the angle between the Y-axis and a line defined by joining the origin to the coordinates (*τ*(*x*), *β*(*x*)) defines the buffering angle. In the 3D space curve, the angle between the Y-plane and a line defined by joining the origin to the coordinates (*σ*(*x*), *τ* (*x*), *β*(*x*)) defines the buffering angle.

Finally, in the same way that the buffering angle can be defined by the buffering parameters, the buffering parameters can be defined by the buffering angle. As described by Schmitt (47), the buffering parameters can be defined trigonometrically using the equations:

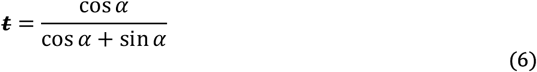

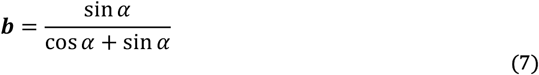

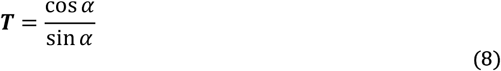

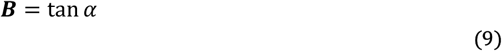

The buffering angle can be calculated from the value of the *t* parameter as follows:

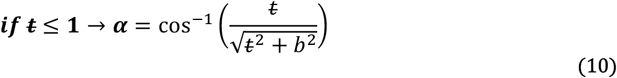

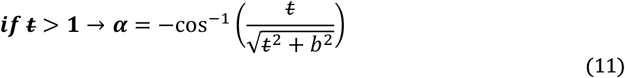

Similarly, the buffering angle can be calculated from the value of the *B* parameter as follows:

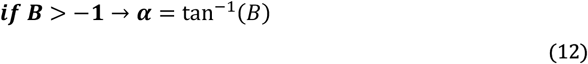

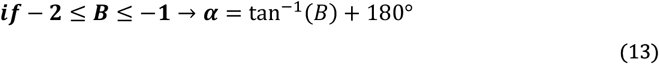

In this way the buffering angle provides a simple, yet powerful, single-value measure that can express the total spectrum of buffering characteristics (from inverted amplification to inversion to moderation to amplification). Furthermore, the angle can be used to calculate the value of *B* (or *T*) so that a fully quantitative measure of buffering on a ratio scale can be provided. In the next section the buffering angle is applied in the context of quantifying genetic buffering.

### Quantifying the buffering effects of gene interactions using the buffering angle

Using the axiomatic framework described above it is now possible to express the expected effects of gene interaction in an intuitive and easily visualizable manner via the buffering angle. To illustrate this further, consider a hypothetical organism with a measurable phenotype, *x*, that varies over time according to the equation *x(t)=x*_*0*_ *rt*^*2*^. Also assume that the wild-type strain exhibits an *r* value of 1/hr, while mutants expressing the ***c***_***1***_ and ***d***_***1***_ alleles exhibit *r* values of 0.7/hr and 0.5/hr, respectively. In this scenario, the buffering parameters of the wild-type reference can be calculated as: *ŧ* =1, *b* =0, *T* is undefined, *B* =0, *α* =0 °. The buffering parameters of the ***c***_***1***_ mutant, on the other hand, can be calculated as: *ŧ* = 0.7, *b*=0 .3, *T* =2.333, *B* =0.429, *α* =23.2°. Likewise, the buffering parameters of the ***d***_***1***_ mutant can be calculated as: *ŧ* = 0.5, *b*=0. 5, *T*=1, *B*=1, *α* =45°.

With respect to the expected phenotype of the double mutant, it is not possible to know *a priori* which neutrality model is applicable (i.e., whether the ***c***_***1***_ and ***d***_***1***_ gene products function in <u>parallel</u> or in <u>series</u>). Thus, the expected buffering parameters for both the parallel and serial models must be calculated. The <u>expected</u> buffering parameters of the ***c***_***1***_***d***_***1***_ double mutant (assuming parallel action) are: *ŧ* = 0.412, *b*=0.588, *T*=0.700, *B*=1.429, *α* =55°. In contrast, the <u>expected</u> buffering parameters of the ***c***_***1***_***d***_***1***_ double mutant (assuming serial action) are: *ŧ* = 0.35, *b*=0.65, *T* =0.538, *B* =1.857, *α* =61.7°.

At this juncture one critical question remains: How can the present axiomatic framework be used to precisely quantify the magnitude of genetic buffering (as defined by Hartwell), and thus the potential for the accumulation of cryptic variation? To answer this question, let us assume that the ***c***_***1***_***d***_***1***_ double mutant is inviable (i.e., ***c***_***1***_ and ***d***_***1***_ are synthetically lethal). The <u>observed</u> buffering parameters of the double mutant strain are thus: *ŧ* =0, *b*=1, *T*=0, *B* is undefined, *α* =90°. For the purposes of this hypothetical scenario, the ***c***_***1***_ allele will also be assumed to encode a true null (i.e., the encoded gene-product is completely non-functional).

In a ***d***^***+***^ genetic background, the expression of the null ***c***_***1***_ allele results in a 23.2° deviation in the buffering angle relative to the wild-type reference (**Figure 7A**). Thus, in a ***d***^***+***^ background there is a 23.2° limit with respect to the expression of phenotypes associated with alleles of ***c*** (which range in functionality from 0% to 100%). However, in a ***d***_***1***_ null background, the possible deviation in the buffering angle is extended to 90°. In other words, in a ***d***^***+***^ background, the possible phenotypic variation resulting from genotypic variation in ***c*** ranges from *B* =0 to *B*=0.429 (i.e., *r* values ranging from 1/hr to 0.7/hr). In a ***d***_***1***_ null background, on the other hand, phenotypic variation resulting from genotypic variation in **c** ranges from *B* =0 to *B* =+∞. Thus, phenotypes corresponding to *B* values greater than 0.429 to +∞ (i.e., *r* <0.7/hr to *r*=0) now become discernable (both to an observer and to natural selection).

**Figure 7.**
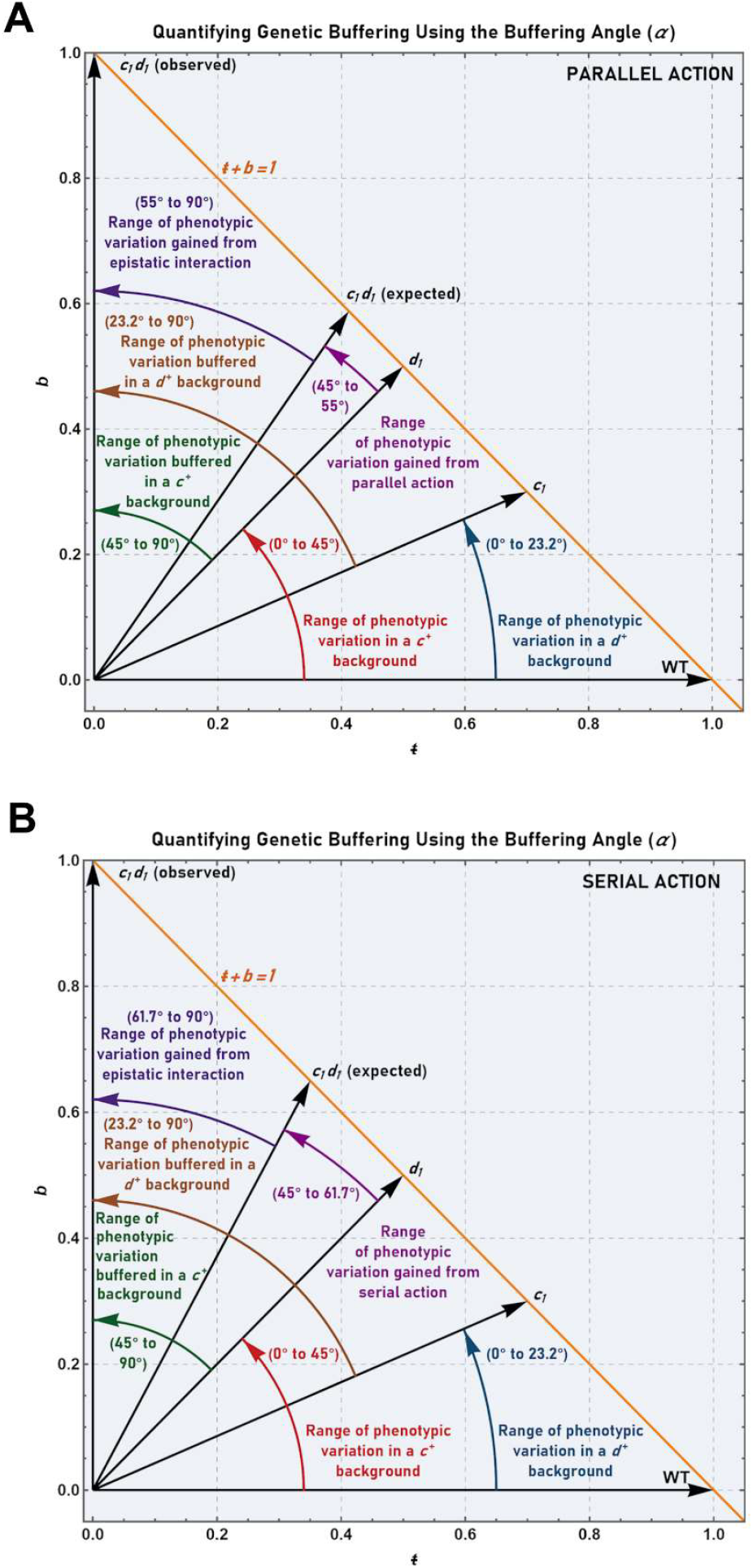
Quantifying genetic buffering using the buffering angle in the context of parallel or serially acting alleles. **(A)** Quantification of the range of phenotypic variation buffered due to genetic interaction between two synthetically lethal and parallel acting alleles (***c***_***1***_ and ***d***_***1***_). For ***c***_***1***_: *ŧ* = 0.7, *b*=0. 3, *T*=2.333, *B* =0.428, and *α* =23.2°. For ***d***_***1***_: *ŧ* = 0.5, *b*=0.5, *T*=1, *B* =1, and *α* =45°. The buffering angles associated with each genotype (black arrows), together with their interpretations, are indicated directly on the plot. The line *b*=1 -*ŧ* is plotted in orange. **(B)** Quantification of the range of phenotypic variation buffered due to genetic interaction between two synthetically lethal and serially acting alleles (***c***_***1***_ and ***d***_***1***_). For ***c***_***1***_: *ŧ* = 0.7, *b*=0.3, *T*=2.333, *B*=0.428, and *α* =23.2°. For ***d***_***1***_: *ŧ* = 0.5, *b*=0.5, *T*=1, *B* =1, and *α* =45°. The buffering angles associated with each genotype (black arrows), together with their interpretations, are indicated directly on the plot. The line *b* =1 - *ŧ* is plotted in orange.

Similarly, in a ***c***^***+***^ background, expression of the ***d***_***1***_ allele (also assumed to be a true null) results in a 45° deviation in the buffering angle relative to the wild-type reference. Thus, in a ***c***^***+***^ background there is a 45° limit with respect to the expression of phenotypes associated with alleles of ***d*** (of varying functionality). However, in a ***c***_***1***_ null background, the possible deviation in the buffering angle is extended to 90°. Thus, in a ***c***^***+***^ background, phenotypic variation caused by genotypic variation in ***d*** ranges from *B* =0 to *B* =1. This contrasts with ***c***_***1***_ null backgrounds, where phenotypic variation caused by genotypic variation in ***d*** ranges from *B*=0 to *B*=+∞. Therefore, phenotypes corresponding to *B* values > 1 to +∞ (i.e., *r* <0.5/hr to *r*=0) become discernable to an observer and to natural selection (**Figure 7A**).

Furthermore, since a 55° deviation was expected (in contrast to the 90° deviation that was observed), 35° of deviation can be cryptically buffered by the genetic relationship between ***c*** and ***d***. In other words, phenotypic variation ranging from *B* =1.429 to *B*=+∞ (i.e., *r* values ranging from 0.412/hr to 0) can effectively be hidden in the presence of a fully functional allele at ***c*** or ***d***. However, at the same time, all or some of this variation may be revealed by subsequent mutations that reduce the functionality of the other gene of the interacting pair. This, of course, would not be possible in the absence of genetic interaction between ***c*** and ***d***. In that case, only phenotypes ranging from *B* =0 to *B* =1.429 would be visible to an observer or natural selection.

To illustrate this further, consider a more extreme example where *α* =0° in both single mutants, but is 90° in the double mutant (the scenario first envisioned by Hartwell). In this instance it is clear that **1)** no phenotypic variation (due to allelic variation in ***c***) is possible in a ***d***^***+***^ genetic background, and **2)** that no phenotypic variation (due to allelic variation in ***d***) is possible in a ***c***^***+***^ genetic background. Thus, genotypic variation in ***c*** is 100% buffered (i.e., completely cryptic) in a ***d***^***+***^ background, and genotypic variation in ***d*** is 100% buffered in a ***c***^***+***^ background.

At the other extreme, consider two mutant null alleles, ***c***_***2***_ and ***d***_***2***_, where *α* is near 90° for each single mutant as well as the double mutant. In this instance, it is clear that **1) *d***^***+***^ has little to no capacity in limiting (i.e., buffering) phenotypic variation arising from allelic variation in ***c***, and **2)** that ***c***^***+***^ has little to no capacity in limiting phenotypic variation arising from allelic variation in ***d***. Thus, genotypic variation in either ***c*** or ***d*** is 100% non-cryptic (i.e., fully discernable to an observer or natural selection). Finally, in intermediate cases (such as the original scenario described above), where phenotypic variation is partially restricted, one can conclude that genotypic variation in either ***c*** or ***d*** is buffered, albeit incompletely, by the other locus of the pair. Most importantly, it is now possible to quantitatively describe the degree of buffering in two ways: **1)** in an intuitive and easily visualizable manner using the buffering angle, or **2)** using a ratio scale – the highest scale possible – based on the *B* parameter (**Figure 7A, B**).

### Genetic interaction scales

In addition to allowing the quantification of buffering action, the current axiomatic framework also provides a simple means by which to classify interactions between alleles at different loci. This is achieved using scales that employ either the *B* parameter (in the case of parallel action) or the *ŧ* parameter (in the case of serial action). To create such a scale, it is first necessary to define a measure of the deviation between **1)** the observed value of *B* or *ŧ* in the double mutant (*B*_*cd*_ or *ŧ* _*cd*_ respectively) and **2)** the expected value of *B* or *ŧ* in the double mutant (*B*_*c*_ + *B*_*d*_ or *ŧ*_*c*_*· ŧ*_*d*_ respectively). This can be accomplished by defining the relative error, *ε*, using the equations below:

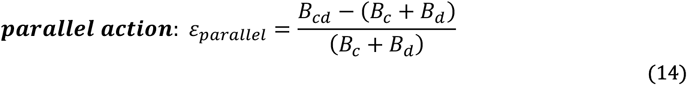

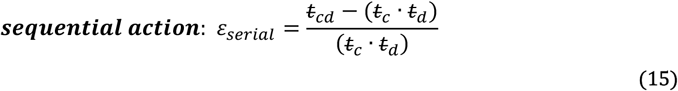

### Parallel action

From inspecting Equation 14 it is evident that *ε*_*parallel*_ evaluates to zero in the absence of genetic interaction between the two alleles (i.e., a score of zero indicates that the phenotypic effects of the expression of the individual alleles at each respective locus are simply <u>additive</u> with respect to the *B* parameter). Positive values of *ε*_*parallel*_, on the other hand, correspond to aggravating interactions where the magnitude of the measured phenotype is more severe than expected. For reference, a value of one on this scale indicates that the observed phenotype is 100%, or two times more severe than expected. A value of two on the scale indicates the observed phenotype is 200%, or three times more severe than expected, and so on. In contrast negative values of *ε*_*parallel*_ correspond to ameliorating interactions where the magnitude of the measured phenotype is less severe than expected. For example, a score of -0.5 would indicate that the double mutant exhibits a 50% decrease in the severity of the phenotype relative to that expected, and a score of -0.9 would indicate a 90% decrease in the severity of the phenotype relative to that expected. A score of -1 indicates that the double mutant exhibits a phenotype that is equivalent to that observed in the wild-type reference. In the context of growth, scores of less than -1 (i.e., scores from -1 to - ∞) indicate that the growth rate of the double mutant exceeds that of the wild-type reference. In contexts other than growth, such scores indicate the effect to be of such magnitude that it exceeds the magnitude of the phenotype exhibited by the wild-type reference.

To aid in judging the effect of such interactions it is possible to define landmarks along the scale described above. For example, an *ε* value corresponding to the case where the value of *B* in the double mutant is equal to that observed in the single mutant exhibiting the weakest phenotype (i.e., the smaller *B* value) can be defined by:

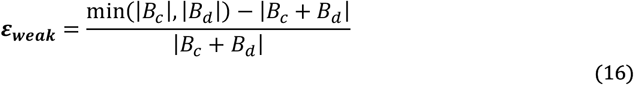

An *ε* value corresponding to the case where the value of *B* in the double mutant is equal to that observed in the single mutant exhibiting the stronger phenotype (i.e., the larger *B* value) can be defined by:

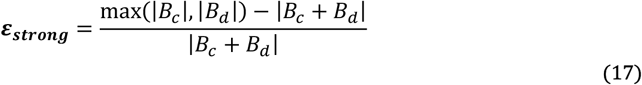

Values of *ε* falling between *ε*_*weak*_ and *ε*_*strong*_ correspond to *B* values intermediate between those observed in the two single mutants. A schematic describing the interaction scale for genes acting in parallel, including the landmarks associated with the scale, is shown in **Figure 8A**.

**Figure 8.**
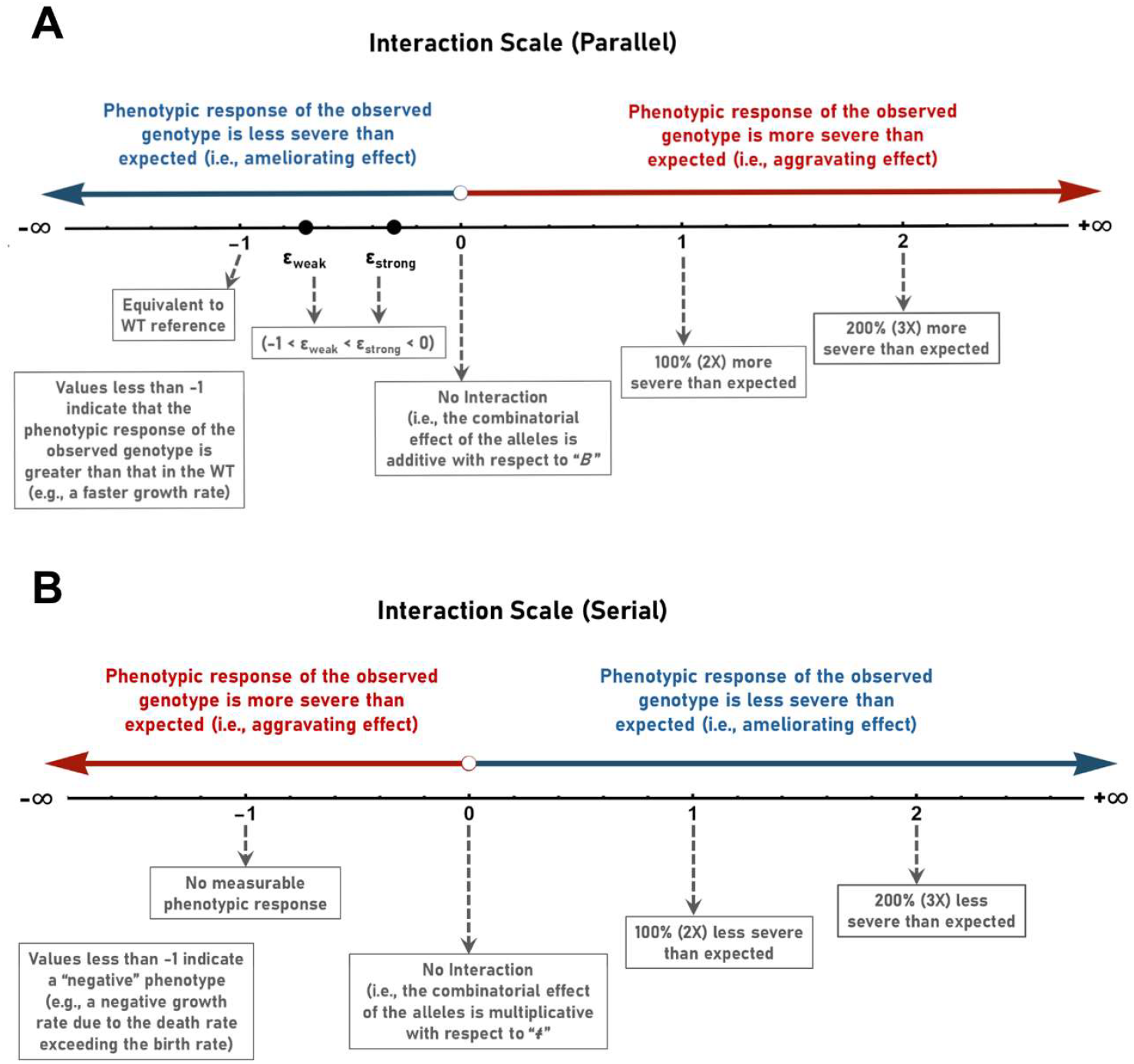
The classification of genetic interactions using scales based on the difference between the observed and expected values of *B* or *t*. (**A**) The strength of interaction between parallel acting genes can be plotted on a scale ranging from -∞ to +∞ based on the relative error between the observed and expected values of the *B* parameter (according to the equation, *ε*_*parallel*_ =(*B*_*cd*_ - *B*_*c*_ -*B*_*d*_)/(*B*_*c*_ +*B*_*d*_). *B*_*cd*_ represents the observed value of *B* in the ***c***_***1***_***d***_***1***_ double mutant. *B*_*c*_ represents the observed value of *B* in the ***c***_***1***_ single mutant. *B*_*d*_ represents the observed value of *B* in the ***d***_***1***_ single mutant. (**B**) The strength of interaction between serially acting genes can be plotted on a scale ranging from - ∞ to +∞ based on the relative error between the observed and expected values of the *ŧ* parameter (according to the equation, *ε*_*serial*_ =(*ŧ*_*cd*_ - *ŧ*_*c*_ · *ŧ*_*d*_)/(*ŧ*_*c*_ · *ŧ*_*d*_). *ŧ*_*cd*_ represents the observed value of *ŧ* in the ***c***_***1***_***d***_***1***_ double mutant. *ŧ*_*c*_ represents the observed value of *ŧ* in the ***c***_***1***_ single mutant. *ŧ*_*d*_ represents the observed value of *ŧ* in the ***d***_***1***_ single mutant.

### Serial action

As revealed by inspection of Equation 15, *ε*_*serial*_ evaluates to zero in the absence of genetic interaction between the two alleles (i.e., a score of zero indicates that the phenotypic effects of the expression of the individual alleles at each respective locus are simply <u>multiplicative</u>). Positive values of *ε*_*serial*_ correspond to ameliorative interactions where the magnitude of the measured phenotype is less severe than expected. For reference, a value of one on this scale indicates that the observed *ŧ* value is 100%, or two times greater than expected. A value of two on the scale indicates the observed *ŧ* value is 200%, or three times greater than expected, and so on. In contrast, negative values of *ε*_*serial*_ correspond to aggravating interactions where the magnitude of the measured phenotype is more severe than expected. For example, a value of - 0.5 indicates that the observed *ŧ* value is half of that expected, and a score of - 0.9 indicates that the observed *t* value is 10% of that expected.

Finally, a score of -1 indicates that the observed *t* value is 0 (i.e., there is no measurable phenotype). A schematic describing the interaction scale for genes acting sequentially is shown in **Figure 8B**.

### Gene interaction diagrams

Lastly, to further aid in the visualization and analysis of genetic interactions, I propose a formal diagrammatic method capable of providing a simple visual representation of the underlying mathematical framework. In the proposed system, genes are represented by nodes. A single gene is represented by a circle labeled with the appropriate gene name or symbol. The circle is filled in when considering a fully functional allele. If a non- or partially functional allele is being considered, then the circle is left unfilled. In the case of an interacting gene pair (e.g., the ***c*** and ***d*** genes) where the alleles being considered function in parallel, a circle bisected by a vertical line is used (with each half of the circle labelled with the appropriate gene name or symbol). When considering the ***c***^***+***^***d***^***+***^ genotype, both halves of the circle are filled in. When considering the ***c***^***-***^***d***^***+***^ genotype, the semi-circle representing ***c***^***-***^ remains unfilled. When considering the ***c***^***+***^***d***^***-***^ genotype, the semi-circle representing ***d***^***-***^ remains unfilled. When considering the ***c***^***-***^***d***^***-***^ genotype, both halves of the circle remain unfilled.

Edges connect nodes to either buffering or transfer compartments (indicated using squares). Edges linking a node to a transfer compartment are labelled with the expected value of the *ŧ* parameter. Similarly, edges linking a node to a buffer compartment are labelled with the expected value of the *b* parameter. Above the square representing each compartment, the expected values of the *b* or *ŧ* parameters are indicated. In the case of parallel action these values are necessarily equal to those attached to each edge. In this way, the summation of values across each horizontal level of the diagram must be equal to 1. In addition, if known, the observed values of the *ŧ* and *b* parameters can be indicated in brackets next to the expected values. An example of such a representation is shown in **Figure 9A**.

**Figure 9.**
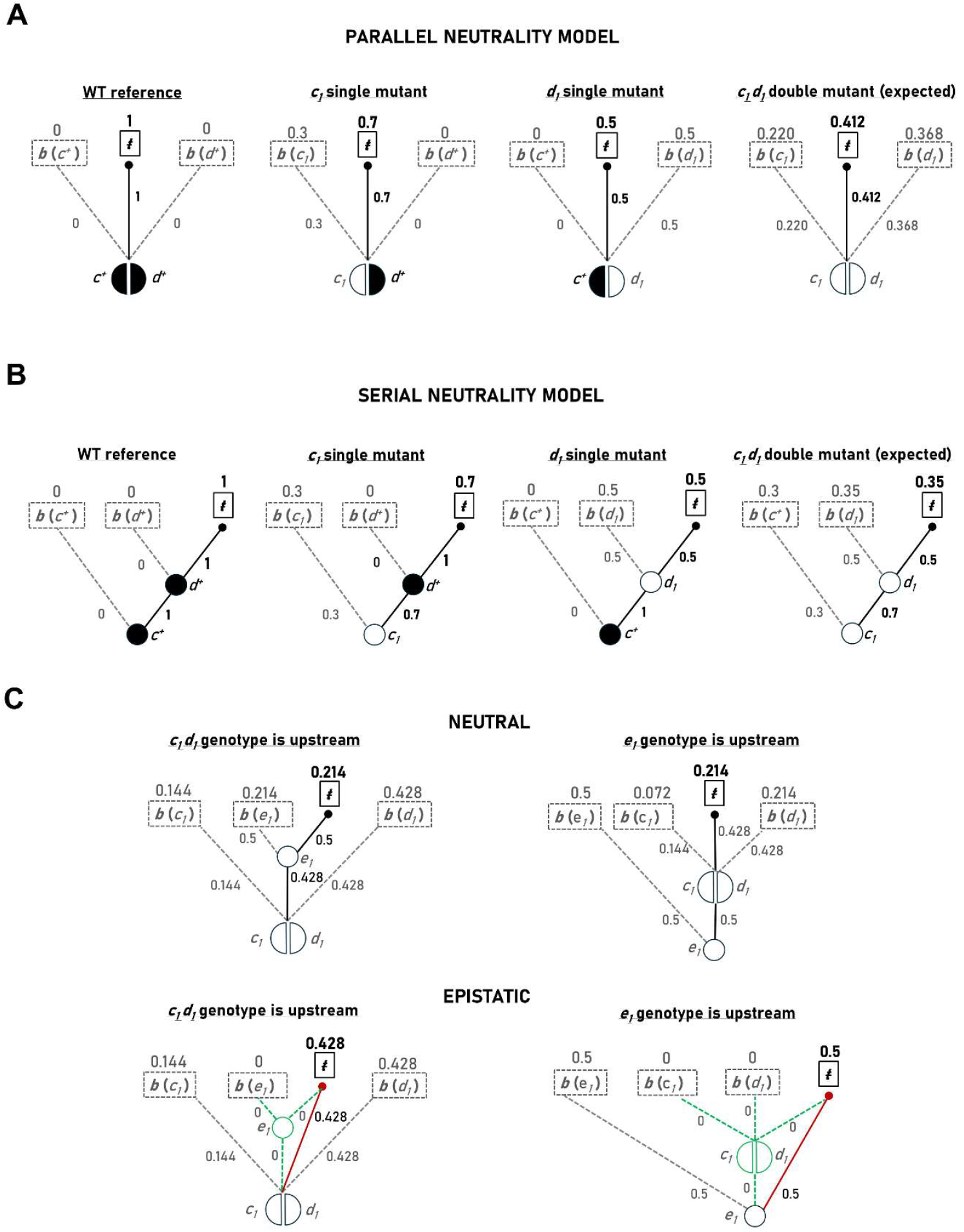
Using genetic interaction diagrams to determine the predicted phenotypic effects of two or more parallel or serially acting alleles. (**A**) Genotypes corresponding to the wild- type reference (*ŧ* = 1, *b*=0, *T* is undefined, *B* =0, *α* =0), the ***c***_***1***_ single mutant (*ŧ* = 0.7, *b*=0.3, *T*=2.333, *B* =0.428, *α* =23.2°), the ***d***_***1***_ single mutant (*t* =0.5, *b*=0.5, *T* =1, *B* =1, *α* =45°), and the ***c***_***1***_***d***_***1***_ double mutant (*ŧ* = 0.412, *b*=0.588, *T* =0.7, *B*=1.429, *α* =55°) are displayed in the scenario where ***c*** and ***d*** act in parallel. Each tree describes a unique genotype. The calculated *ŧ* parameter for each genotype is shown in bold above the “*ŧ*” compartment (bold rectangle). The calculated *b* parameters are shown in grey above the respective “*b"* compartments (grey rectangles). The sum of the *ŧ* and *b* values across any horizontal layer of a tree must equal one. (**B**) Genotypes corresponding to the wild-type reference (*ŧ* = 1, *b*=0, *T* is undefined, *B* =0, *α* =0), the ***c***_***1***_ single mutant (*ŧ* = 0.7, *b*=0.3, *T* =2.333, *B* =0.428, *α* =23.2°), the ***d***_***1***_ single mutant (*ŧ* = 0.5, *b*=0.5, *T*=1, *B* =1, *α* =45°), and the ***c***_***1***_***d***_***1***_ double mutant (*ŧ* =0.35, *b*=0.65, *T*=0.538, *B* =1.857, *α* =61.7°) are displayed in the scenario where ***c*** and ***d*** act sequentially. Each tree describes a unique genotype. The calculated *ŧ* parameter for each genotype is shown in bold above the “*ŧ*” compartment (bold rectangle). The calculated *b* parameters are shown in grey above the respective “*b"* compartments (grey rectangles). The sum of the *ŧ* and *b* values across any horizontal layer of a tree must equal one. Values of *t* or *b* along any vertical path are multiplied to calculate the value of *ŧ* or *b* passed to the relevant transfer or buffer compartment. (**C**) Genetic interaction diagrams describing a possible three-gene scenario. The top panel describes non-epistatic (i.e., neutral) interactions between ***c***_***1***_ (*ŧ* =0.75, *b*=0.25, *T* =3, *B* =1/3, *α* =18.4°), ***d***_***1***_ (*ŧ* =0.5, *b*=0.5, *T*=1, *B* =1, *α* =45°), and ***e***_***1***_ (*ŧ* = 0.5, *b*=0.5, *T*=1, *B*=1, *α* =45°), where ***c***_***1***_ and ***d***_***1***_ act in parallel and ***e***_***1***_ is either downstream (left) or upstream (right) in the pathway. The bottom panel describes an example of an epistatic (i.e., non-neutral) interaction between the alleles, where ***c***_***1***_ and ***d***_***1***_ act in parallel and ***e***_***1***_ is either downstream (left) or upstream (right) in the pathway.

In scenarios involving serial action, edges are not only permitted to link nodes to buffer or transfer compartments, but are also permitted to link nodes to other nodes. Edges connecting nodes are labelled with the value of the *ŧ* parameter. In cases where gene ***c*** is linked by an edge to gene ***d***, which in turn is linked to a transfer compartment, the individual *ŧ* values are multiplied and the expected value placed above the transfer compartment. Thus, *ŧ* values along any vertical path are multiplied. As before, edges linking a node to a buffer compartment are labelled with the expected value of the *b* parameter. Like the scenario involving parallel acting genes, the summation of values across each horizontal level of the diagram must again equal one. In addition, if known, the observed values of the *ŧ* and *b* parameters can be indicated in brackets next to the expected values. An example of such a representation is shown in **Figure 9B**.

Furthermore, it is important to note that these same rules can be applied to more complex scenarios involving three or more genes, as well as those involving both serial and parallel action. A gene interaction diagram describing such a three gene scenario is presented in **Figure 9C**. Since each “tree” represents a distinct genotype and provides both a visual and quantitative description of the nature of the interaction being analyzed, the proposed diagrammatic system is sufficiently flexible and scalable to accommodate complex pathways or regulatory modules that would otherwise be beyond human intuition. It is thus hoped that this approach will prove useful in the systematic analysis of complex interactions involving multiple genes.

## Discussion

### A quantitative terminology for describing gene interaction

In the preceding sections, a formal axiomatic framework for the quantitative analysis of genetic buffering was presented (using the work of Schmitt (47) as a foundation). Simple extensions of this framework provide a means to quantitatively describe the phenotypic effects of individual mutant alleles as well as interactions between alleles in a formal, general, and mathematical way. Thus, the qualitative parlance often used to describe gene interaction (e.g., positive, negative, synergistic, antagonistic, ameliorative, aggravating, diminishing, compensatory, reinforcing, masking, suppressing) can now be replaced by a quantitative description based on the deviation of *B*_*observed*_ from *B*_*expected*_ (in the case of parallel action) or on the deviation of *ŧ*_*observed*_ from *ŧ*_*expected*_ (in the case of serial action). This is a welcome advance given that such qualitative terms have been the source of much confusion within the discipline. Importantly, the methodology also allows the concept of genetic buffering (as conceptualized by Hartwell) to be defined in a precise and quantitative manner as a specific case within the extended framework. Finally, a formal set of rules for quantifying and comparing gene interactions, as well as for creating intuitive visual representations of complex gene interaction scenarios, has been devised to assist in their systematic analysis.

The broader implications of this work with respect to the definition of epistasis and the quantitative analysis of genetic phenomena are further expanded upon below.

### The axiomatic basis of the approach

The mathematical framework herein described rests on three axioms:

1) The phenotypic effect of an allele can be represented by a partitioning process defined by **i)** the transfer function, *τ(x)*, **ii)** the buffer function, *β(x)*, and **iii)** the sigma function, *σ(x)*, where *σ(x) = τ(x) + β(x)*.
2) The state of the buffered system can be described in its entirety by one of the five previously defined buffering parameters, *ŧ, b, T, B*, or *α*.
3) Interaction between two mutant alleles at different loci can be quantitatively assessed by comparing the observed phenotypic effect in the double mutant to that expected upon applying one of two mutually exclusive neutrality models, the “parallel” neutrality model or the “serial” neutrality model.

### Notes on the first axiom

The transfer function, *τ(x)*, describes the quantity of interest (i.e., the aspect being measured) as observed in a mutant genotype relative to a reference genotype. The choice of reference genotype is arbitrary, but one expressing the wild-type phenotype makes the derived numerical results the most intuitively interpretable (notwithstanding the difficulty in defining the meaning of “wild type” in the context of natural populations). The sigma function, *σ(x)*, describes the quantity of interest as observed in the reference genotype relative to itself, hence *σ(x)=x*. The buffer function, *β(x)*, is simply the difference between *σ(x)* and *τ(x)* and describes the quantity of interest as inferred to exist within an abstract buffering compartment (i.e., an imagined compartment containing the phenotypic output <u>not</u> observed in the mutant due to the buffering action of the allele).

Up to this point, the derivation of these functions has been described intuitively using the analogy of fluid filled vessels and has not been examined in a genetical context. This aspect of the procedure is expanded upon below.

### The buffering functions in a genetical context

Deriving the sigma function first requires the creation of a model of the quantity of interest, *x*, where *x* represents the attribute being observed and measured. Using this model, a function describing the quantity in relation to time, *x*(*t*), can be generated. In the case of exponential growth, the quantity of interest is simply the number of cells in the population (indicated with the variable, *N*) and the model describing exponential growth is the ODE, *dN/dt=rN*. To generate *N* as a function of time, *dN/dt=rN* is integrated to give *N(t)=N*_*0*_ *e*^*rt*^. Importantly, *N(t)=N*_*0*_ *e*^*rt*^ can itself be integrated to evaluate the “accumulated” phenotype over a given observation period. For example, if one assumes a 10-hour observation period, then the subsequent integration of *N(t)* itself (from *t=* 0 hours to *t*=10 hours) gives the <u>definite</u> integral:

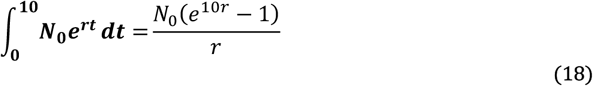

This, in essence, evaluates the “cumulative” or “integrated” phenotypic expression over the period of observation as a real number in units of cell·hours (or more generally, the units of the measured trait sustained per unit time). The use of cell·hours as units in this context is analogous to the way that kilowatt·hours are used to describe the delivery of energy over time, or how person·hours are used to describe the total amount of work performed by a group of individuals over a given interval of time. In this way “cell·hours” represents the fundamental unit of measurement with which to describe microbial growth. Similarly, the fundamental units with which to describe a given phenotype are the units of <u>trait</u> measurement multiplied by the units of <u>time</u> measurement.

It is also possible to define the <u>indefinite</u> integral of *N(t)*,

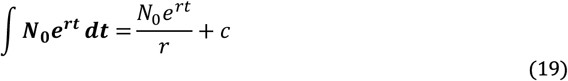

where *c* represents the constant of integration. This function is most aptly referred to as the “plenary” function since it represents the indefinite integrated expression of the observable phenotype. Thus, upon examining this process through the lens of the associated time derivatives, the function *x(t)* is simply the first derivative of the plenary function.

In any event, from a genetical context, the range of *x(t)* over the observation period defines the values of the quantity of interest witnessed in the wild type and thus provides the frame of reference needed to compare mutant alleles. This is achieved mathematically by defining the transfer function, *τ(x)*, which describes the phenotypic output observed in a mutant versus the phenotypic output that is possible (as determined by the observation of the wild-type reference). It is for this reason that *σ(x)=x* (i.e., the phenotypic output in the wild type is equal to that observed in the reference by definition). Lastly, using this same logic, the “unrealized” phenotypic output in the mutant can simply be defined by the buffer function as *β(x)= σ(x)- τ(x)*.

It should also be noted that an approach similar to that described above could be taken to define functions describing the effects of environmental (as opposed to genetical) perturbations on the phenotypic output of a reference genotype. In this way, the effects of both genetical and environmental factors could be incorporated into a single, common mathematical framework. This will be explored further in a subsequent manuscript.

### Time dilation/compression

It is also useful to note that the differences between wild-type and mutant genotypes can be viewed through the lens of the time dimension as well as the phenotypic dimension. For example, the consequences of a two-fold reduction in a measured phenotype could be viewed using either time or phenotypic response as the point of reference. This is to say, for a phenotype varying linearly with time, one could not only view the mutant as achieving half the phenotypic output of the wild type over the <u>same length of time</u>, but one could view the mutant as taking twice as long as the wild type in achieving <u>the same phenotypic output</u>. For phenotypes varying linearly with time, the time dilation (*ŧ*<1) or compression (*ŧ*>1) experienced in the mutant can be defined *by t*_*mutant*_*=t*_*WT*_*/ŧ*.

More generally, for phenotypes varying non-linearly with time, *t*_*mutant*_ can be defined by:

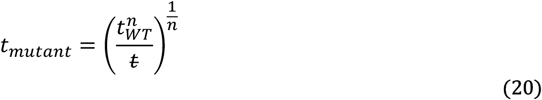

where *n* is the degree of *t* (time) in the function *x(t)* describing the quantity of interest.

### Notes on the second axiom

The state of any buffered system can be described in its entirety by one of the five buffering parameters: *ŧ, b, T, B*, and *α*, where *ŧ* and *b* represent the respective changes in the transfer and buffer compartments relative to the total change, and *T* and *B* represent the change in either of the two compartments relative to the other. The buffering angle, *α*, on the other hand, is a trigonometric representation of the system that provides two distinct advantages: **1)** the description of the system in the absence of discontinuities brought about by division by zero, and **2)** the ability to graphically display its characteristics in an intuitive, yet mathematically precise manner.

As noted by Schmitt (47), the analysis of buffered systems using this paradigm is somewhat analogous to the quantitation of chance events using either <u>probabilities</u> (comparable to the description of buffered systems using the *ŧ* or *b* parameters) or <u>odds</u> (comparable to the description of buffered systems using the *T* or *B* parameters). The former allows the description of the system using a relative scale (normalized to 1), whereas the latter allows the description of the system using a ratio scale (i.e., a scale with a non-arbitrary zero and equal intervals between neighboring points). Ratio scales are particularly useful as they allow for the addition, subtraction, multiplication, and division of values, as well as their statistical description (arithmetic mean, median, mode, range, variance, standard deviation). For these reasons ratio scales are considered the highest scale possible (49). From a genetical standpoint, the current framework thus allows the full quantitative analysis of gene interactions across organisms using a common methodology or “toolbox".

Using this common toolbox, any phenotypic effect or gene interaction can be described provided a quantitative methodology capable of reliably gauging the phenotype of interest exists. This is to say, for any phenotype of interest, an operationally defined measurement process that is both reproducible and valid must be available (51). Importantly, the parameters as herein defined are capable of coping with scenarios involving negative *b* or *ŧ* values. For example, growth models that incorporate a death/mortality rate in addition to a birth/proliferation rate (making negative growth a possibility) necessitate the use of negative *ŧ* values which are easily accommodated mathematically within the framework.

### Notes on the third axiom

In determining the expected phenotypic effect of the simultaneous expression of two mutant alleles at distinct loci, one of two mutually exclusive neutrality models must be chosen: the “parallel” neutrality model or the “serial” neutrality model. In the absence of data regarding the biochemical or molecular function of the respective gene-products (that could be used to distinguish between their parallel or serial action), it is not possible to know *a priori* which of these models should be applied. In cases involving parallel action, the expected phenotypic effect is calculated through the addition of the *B* values determined for each of the single mutants. In cases involving serial action, the expected phenotypic effect is calculated by multiplying the individual *ŧ* values determined for each of the single mutants. In either case, the value of any of the other buffering parameters corresponding to the calculated *B*_*expected*_ or *ŧ*_*expected*_ value can be determined if desired. Furthermore, the relative error of *B*_*observed*_ or *ŧ*_*observed*_ can be used to generate scales that can be used to quantitatively classify a genetic interaction.

Using this approach, it becomes clear that **1)** genetic buffering (in the Hartwellian sense) is simply a specific case where *B*_*observed*_>> *B*_*expected*_ and **2)** that the magnitude of the difference between *B*_*observed*_ and *B*_*expected*_ indicates the increase in the range of phenotypic variation that may be accessible to natural selection. Given that the parallel neutrality model has not been previously considered, it would be of some interest to re-analyze past genetic interaction data with this model in mind.

### The definition of epistasis

Based upon the principles described above, it becomes possible to provide a precise and quantitative definition of epistasis. Speaking in purely mathematical terms, epistasis can now be defined as the case where either *B*_*cd*_ ≠ *B*_*c*_ *+ B*_*d*_ (for parallel acting alleles), or where *ŧ*_*cd*_ ≠ *ŧ*_*c*_*·ŧ*_*d*_ (for serially acting alleles). It is interesting to note that this definition provides an underlying mathematical rigor to the most common qualitative definition (based on the “Fisher” paradigm) that is in current use – **an interaction between alleles at different loci resulting in a phenotype distinct from that expected under the assumption of their independent action** (15).

In contrast, if *B*_*cd*_ is <u>not</u> significantly different from *B*_*c*_ + *B*_*d*_, or if *ŧ*_*cd*_ is not significantly different from *ŧ*_*c*_·*ŧ*_*d*_, then the relationship between the two alleles can be classified as being non-epistatic. Importantly, such non-epistatic relationships can be further sub-divided into those that are additive (in the case of parallel action) and those that are multiplicative (in the case of serial action). Thus, three classes of relationship exist between alleles at different loci (with respect to their effect on a given phenotype): **1)** additive, **2)** multiplicative, and **3)** epistatic. Furthermore, systems of interacting alleles (involving all three classes of relationship) can now easily be described in a formal an axiomatic manner using gene interaction diagrams. Many excellent reviews on the definition of epistasis are available (14, 15, 42, 52–54) and would be useful to refer to in comparing the viewpoint presented here with other alternatives.

Lastly, it is of particular interest to note the work of Zuk and Lander (22), who suggest that the “missing heritability” arises from an over-estimate of the total (narrow sense) heritability. Furthermore, the authors suggest that this over-estimate results from the implicit assumption that gene interactions are <u>not</u> involved in the expression of the trait being observed. This raises two significant questions: **1)** Do genetic buffering relationships explain all (or some) of this underestimate? and **2)** Can the current framework be used to quantify the effects of such interactions with respect to the extent of cryptic variation (at least in experimentally controlled populations)? These questions go beyond the scope of the current discussion and will be explored further in a subsequent manuscript. In any event, it is hoped that the current framework will provide a general model of epistasis that is broadly applicable to all living organisms and that is capable of quantitatively communicating the phenotypic effects of individual alleles and their interactions in a rigorous and mathematical manner.

## Acknowledgments

The author acknowledges the support of the Natural Sciences and Engineering Research Council of Canada (funding reference number, R4029A03).

## Notes

### Competing Interest Statement

The authors have declared no competing interest.

